# Low diversity and microdiversity of comammox bacteria in wastewater systems suggests wastewater-specific adaptation within the *Ca*. Nitrospira nitrosa cluster

**DOI:** 10.1101/2022.06.11.495745

**Authors:** Irmarie Cotto, Katherine J. Vilardi, Linxuan Huo, Emily C. Fogarty, Wendell Khunjar, Christopher Wilson, Haydee De Clippeleir, Kevin Gilmore, Erika Bailey, Sebastian Lücker, Ameet J. Pinto

## Abstract

Studies have found *Ca*. Nitrospira nitrosa-like bacteria to be the principal or sole comammox bacteria in nitrogen removal systems for wastewater treatment. In contrast, multiple populations of strict ammonia and nitrite oxidizers co-exist in similar systems. This apparent lack of diversity is surprising and could impact the feasibility of leveraging comammox bacteria for nitrogen removal. We used full-length 16S rRNA gene sequencing and genome-resolved metagenomics to compare population-level (i.e., species) diversity of comammox bacteria with that of strict nitrifiers in full-scale wastewater treatment systems and assess whether these observations were consistent or diverged at the strain-level. Full-length 16S rRNA gene sequencing indicated that while *Nitrosomonas*-like bacteria exhibited higher population-level diversity, the effective microdiversity of most *Nitrospira*-like bacteria were comparatively higher except for one *Nitrospira* Lineage II population. Comammox bacterial metagenome assembled genomes (MAGs) were associated with *Ca*. Nitrospira nitrosa. The average amino acid identity between comammox bacterial MAGs (93% ± 3) across systems was significantly higher than that of the *Nitrosomonas*-like ammonia oxidizers (73%±8) and the *Nitrospira*-like nitrite oxidizer MAGs (75%±13), suggesting that the same comammox population was detected in all systems. Comammox bacteria and some ammonia oxidizers MAGs were significantly less microdiverse than most ammonia and nitrite oxidizers. Interestingly, strain-resolved analysis also indicates that different nitrogen removal systems harbor different comammox bacterial strains within the *Ca*. Nitrospira nitrosa cluster. These results suggest that comammox bacteria associated with *Ca*. Nitrospira nitrosa have low species- and strain-level diversity in nitrogen removal systems and may thus harbor specific adaptations to the wastewater ecosystem.

## Introduction

Aerobic nitrification processes for nitrogen removal from wastewater are largely centered around biotransformation and growth kinetics of strict ammonia oxidizing bacteria (AOB) and nitrite oxidizing bacteria (NOB) (1). Since their discovery (2–5), several studies have detected comammox bacteria in a wide range of environmental and engineered systems (2–9), including nitrogen removal processes in laboratory and full-scale wastewater treatment bioreactors (9–20). Compared to oligotrophic engineered systems such as drinking water (9,21) and even tertiary treatment systems at full-scale wastewater treatment plants (WWTPs) (16) with low nitrogen concentrations, most studies have identified *Candidatus* Nitrospira nitrosa-like comammox bacteria as the principal or sole comammox bacteria in laboratory or full-scale secondary treatment systems (11,22,23). This apparent lack of diversity of comammox bacteria in secondary treatment processes is not only surprising but has the potential to impact treatment strategies centered around comammox bacteria (24,25) as functional outcomes reliant on low diversity communities may be less resilient to environmental fluctuations and perturbations.

*Nitrosomonas*-like AOB and *Nitrospira*-like NOB have been studied extensively in wastewater treatment systems and studies often report the coexistence of multiple populations (26–28) likely occupying different ecological niches (i.e., adapted to different conditions) within a complex community. Functional redundancy of multiple co-exiting populations can confer stability to microbial ecosystems ensuring the long-term persistence (29,30). Despite the apparent low population diversity of comammox bacteria compared to strict nitrifiers, they have exhibited remarkable stability in secondary treatment systems (11,15,31). One plausible reason for their temporal persistence could be that comammox bacteria may harbor intra-population (i.e., strain) level diversity as compared to population level (i.e., species). Previous literature has demonstrated that microdiversity within populations can not only allow them to adapt to rapidly changing conditions, but also contribute to their persistence (7,32). Further, small variations in metabolic capacity may underpin metabolic diversity and thus enable the co-existence of multiple closely related strains within a single population. *Nitrospira*-like bacteria have been shown to harbor high levels of microdiversity in drinking water (33,34) and wastewater (35) systems. Moreover, studies have reported that closely related co-existing *Nitrospira* strains can exhibit differences in substrate affinities, and utilization rates (35). Since, both levels of diversity (i.e., population and intra-population) have been associated with microbial persistence and ecosystem stability, it is plausible that the persistence of comammox bacteria may be associated with higher levels of microdiversity; this would be consistent with observations for *Nitrospira-like* bacteria (35).

To test this hypothesis, we used a combination of full-length 16S rRNA gene sequencing and hybrid assembly and binning approaches to systematically explore the population- and strain-level diversity of nitrifiers in three full-scale nitrogen removal systems with different process configurations. While 16S rRNA gene sequencing cannot provide information on genome content or metabolic potential, the single nucleotide resolution across the full length 16S rRNA gene can help determine differences in (micro)diversity among nitrifying populations (35,36). Nevertheless, 16S rRNA gene sequences cannot be used to reliably distinguish comammox bacteria from strict *Nitrospira*-NOB. One approach to differentiate between the different guilds within *Nitrospira* bacteria and to analyze their diversity at high resolution is to leverage a genome resolved approach to obtain high quality metagenome assembled genomes (MAGs). Various methods exist to quantify diversity (e.g., average nucleotide identity [ANI] and average amino acid identity [AAI]) and microdiversity (e.g., average nucleotide identity from reads and nucleotide diversity from single nucleotide polymorphisms [SNPs]) within populations. Nevertheless, their applicability depends on the quality of MAGs which can be impacted when relying on short read sequencing (e.g., Illumina) in the presence of closely related strains (37); long-read sequencing on the Nanopore platform can help mitigate this constraint (38,39). However, this requires high levels of coverage to obtain polished consensus sequences with lower error rates compared to the raw data (37). Here, we utilize a hybrid metagenomic assembly approach, including long- and short-read data in an effort to assemble high quality MAGs for subsequent microdiversity analyses (37).

Thus, the overall objective of this study was to test the hypothesis that the widespread distribution and persistence of nearly identical comammox bacterial populations in multiple secondary wastewater treatment systems was due to the co-existence of multiple strains (i.e., high microdiversity). To accomplish this, we used a combination of full-length 16S rRNA gene sequencing and hybrid metagenome assembly and binning approach to systematically explore the population and strain level diversity of nitrifiers in three full-scale nitrogen removal systems with different process configurations. In doing so, we also aimed to assess factors that may influence the inter- and intra-population diversity of co-existing nitrifiers.

## 2.0 Materials and Methods

### 2.1 Samples selection and processing

Samples were selected from three nitrogen removal systems with different process configurations (i.e., sequencing batch reactor (SBR), Integrated Fixed Film Activated Sludge (IFAS), and Bardenpho 4-stages activated sludge system) with high concentrations of comammox bacteria in our previous study (31). From this sample archive collected in 2017-2018, a total of thirty-three, twenty, and six samples, respectively, were selected for full-length 16S rRNA gene sequencing on the PacBio Sequel IIe platform, and short and long read metagenomic sequencing on the Illumina NovaSeq and Nanopore MinION platforms, respectively (Table 1). Sample collection and processing (including DNA extraction) and process data collection were described previously (31) and are outlined in the supplementary text.

**Table 1:**
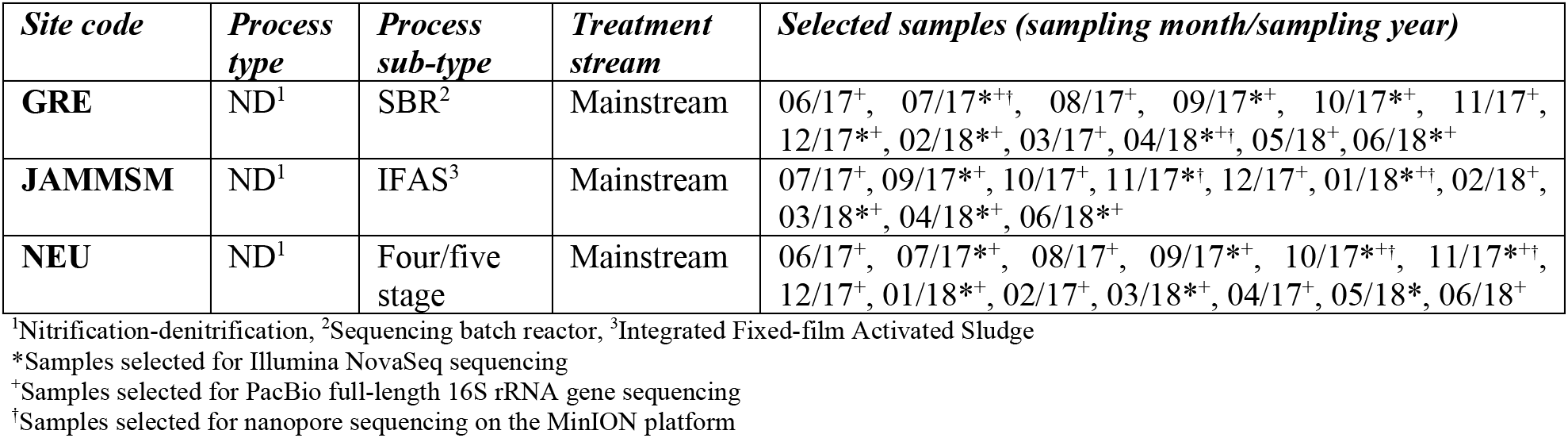
Overview of system process type and sub-type, operational scale and samples included in this study and sample-specific sequencing strategy.

### 2.2 PacBio full-length 16S rRNA gene sequencing and data analysis

Sample DNA extracts (Table 1) were sent for full-length 16S rRNA gene sequencing at the Roy J. Carver Biotechnology Center (Sequencing Core, University of Illinois Urbana-Champaign). The full-length 16S rRNA gene amplicons were generated with universal barcoded primers (27F and 1492R)(40), PCR products were subject to library preparation and sequenced on the PacBio Sequel IIe using the circular consensus sequencing (CCS) mode. Raw reads were demultiplexed and CCS analyses were performed to obtained consensus reads. Further details on PCR, library preparation, and sequencing are provided in the supplementary text. Downstream data processing was performed on a system-by-system basis using DADA2 v1.19.2 (41) in R v4.0.4. using the sample inference method for full-length 16S rRNA gene with single-nucleotide resolution (42). Table S1a summarizes the reads per samples at different stages of data processing in DADA2. ASVs with ≥100 reads in any of the three systems (~>0.01% relative abundance) were clustered into operational taxonomy units (OTUs) using the function ‘IdClusters’ in DECIPHER v2.16.1. (43) at 98.7% sequence identity, a previously recommended threshold for clustering full-length 16S rRNA gene sequences at the species level (44,45). The most abundant ASV within each OTU was used as representative sequence and its taxonomic affiliation was used as the consensus taxonomy for the OTU. Additional details on ASV to OTU data processing are provided in the supplementary text. Representative ASV sequence from all OTUs were aligned with MUSCLE v3.8.1551 (46), and the phylogenetic tree was constructed and visualized using IQ-TREE v2.0.3 (47) and iTOL v2.1.7 (48), respectively. Principal Coordinates Analysis (PCoA) was performed with the weighted and unweighted UniFrac distance metric (49) using the ‘ordinate’ function of phyloseq v1.32.0 and the ‘plot_ordination’ function of ggplot2 v3.3.5 (50).

ASVs classified at the genus level as *Nitrospira* and *Nitrosomonas* (the only two nitrifying genera detected in this study) were extracted from the rarefied ASV table (rarefied to the sample with lowest read count) and reference 16S rRNA gene sequences for the two genera were obtained from the SILVA SSU 138.1 database (Tables S2 and S3). Comammox bacterial 16S rRNA genes were extracted from references genomes downloaded from NCBI (Table S4). MUSCLE v3.8.1551 was used to align ASVs sequences from each genus with their respective references, and a maximum likelihood phylogenetic tree of each species was generated using IQ-TREE v2.0.3 and visualized in iTOL v2.1.7. To assess the population diversity (i.e., genus level) the Shannon diversity was calculated per sample from the relative abundance of the OTUs classified as the same genus (e.g., *Nitrospira)* as the exponential of the Shannon index. To estimate intra-population diversity, the effective microdiversity of each OTU was calculated on a per sample basis from the relative abundance of its ASVs as the exponential of the Shannon index, which is analogous to the effective number of strains (i.e., ASVs) within a population (i.e., OTU) (36).

### 2.3 Illumina and Oxford Nanopore sequencing, co-assembly and hybrid assembly

Seven samples from GRE and NEU and six samples from JAMMSM were sent for sequencing on the Illumina NovaSeq 6000 platform, and two samples from each system on the Oxford Nanopore Technologies GridIONx5 (Table 1) to the Roy J. Carver Biotechnology Center at the University of Illinois Urbana-Champaign Sequencing Core. Library preparation and sequencing details are provided in the supplementary text. These runs resulted in 1.78 billion paired-end short reads (2×150 nt reads) and 45.3 Gbps of long read data (Table S1b and c). Raw short reads were filtered using fastp v0.20.0 (51) and the Univec database was used to remove contamination from the filtered reads as previously described (31). The resulting bam files were sorted using SAMtools v1.9 (52) and converted into fastq files with bedtools v2.29.0 (53). All reads from the same system were co-assembled into contigs using metaSpades v3.13.0 (54). Co-assemblies were performed with kmer sizes of 21, 33, 55 and 77. Contigs smaller than 500 bp were removed from the co-assemblies using the Anvi’o v6.1 command ‘anvi-script-reformat-fasta’ (55). The reformatted assembly fasta files were indexed with bwa index v0.7.17 (56) and the paired end reads from each metagenome were mapped to the respective co-assembly using bwa mem v0.7.17. The resulting sam files were converted to bam files using ‘samtools view -F 4 -bhS’ to retain only mapped reads. Hybrid metagenomic assemblies were performed using OPERA-MS v0.9.0., which combines the advantages of short and long-read technologies to improve genome assemblies (37). The OPERA-MS inputs were short-read metagenomic assemblies to provide a good representation of the sequences in the metagenome, and long and short reads to identify connections between the contigs and obtain contiguous assemblies with low base-pair error. OPERA-MS was executed with the flags --no-ref-clustering and --long-read-mapper minimap2. Both sets of co-assemblies were evaluated using QUAST v5.0.2 (57) (Table S5).

### 2.4 Recovery, annotation, refining and dereplication of metagenome assembled genomes (MAGs)

Binning was performed separately, with co-assemblies from Illumina short reads only and hybrid assemblies from each system using MetaBAT2 v2.12.1 (58), CONCOCT v1.1.0 (59), and MaxBin2 v2.2.7 (60) using contigs greater than 2000 bp. Pilon v1.23 (61) was used for polishing to improve the draft bins obtained from the hybrid assemblies. The quality and taxonomy of the resulting bins were determined with CheckM v1.1.2 (62) and the Genome Taxonomy Database Toolkit (GTDB-Tk 1.1.1, database release r86 v3) (63), respectively. Bins were subject to gene calling using Prodigal v2.6.3 (64) and gene annotation against the KEGG database (65) using kofamscan v1.2.0 (66). Only bins that were taxonomically assigned to known nitrifying genera or those containing genes associated with nitrification (i.e., *amoA* [KO number K10944], *amoB*[K10945], *amoC* [K10946], *hao* [K10535], *nxrA* [K00370], *nxrB* [K00371]) were retained for manual refinement with Anvi’o v6.1. DAStool v1.1.2 (67) was used to combine and curate the refined bins from the three binning methods and generate a non-redundant set of bins from each co-assembly (i.e., one set of bins per system for the short-read only and for the hybrid assemblies). In total, 43 nitrifiers bins from the short-read co-assemblies and 30 from the hybrid assemblies were obtained. These bins were de-replicated using drep v2.5.4 at 95% ANI with completeness and contamination thresholds set to 50% and 10%, respectively. This resulted in 44 nitrifier MAGs with completeness and redundancy estimates higher than 50% and lower than 10%, respectively, of which 21 contained genes associated with nitrification (i.e., *amoA*, *amoB*, *amoC*, *hao*, *nxrA*, and/or *nxrB*).

### 2.5 Taxonomy aware re-assembly using nitrifying bacterial MAGs

We performed re-assemblies from reads mapped to the 44 nitrifying bacterial MAGs to improve MAG quality. First, contigs associated with *Nitrospira* (including comammox) and *Nitrosomonas*MAGs from each system (i.e., GRE, JAMMSM, NEU) were collated into a single fasta file per system. The resulting fasta files were indexed using bwa index v0.7.17 and short reads from the respective system were mapped using bwa mem v0.7.17. The resultant bam file was split into *Nitrospira* and *Nitrosomonas* specific bam files per system and respective fastq files were generated per using samtools fastq. Long reads were also recruited by mapping the reads from Nanopore sequencing to the fasta files with ‘bwa mem -x ont2d’. Short reads mapping to the contigs of the MAGs classified as *Nitrospira* and *Nitrosomonas* were re-assembled using metaSpades v3.13.0 with kmers 21, 33, 55 and 77 on a system-by-system basis. The new genus-specific assemblies and fastq files of mapped long reads were used as input for OPERA-MS to perform hybrid metagenomic re-assemblies (n=9). Quality assessment, binning, taxonomy annotation, manual refining and gene calling were performed as described before. This resulted in 36 nitrifier MAGs with completeness and redundancy estimates higher than 50% and lower than 10%, respectively, of which 24 contained genes associated with nitrification (*amoA*, *amoB*, *amoC*, *hao*, *nxrA*, and/or *nxrB*). The abundance of each MAG per sample (reads per kilobase million, RPKM) and proportion of genome covered was calculated with coverM (68). Four low abundant MAGs (3 AOB and 1 *Nitrospira*-NOB) with less than 50% genome coverage in any of the samples were removed from subsequent analyses. The entire workflow for assembly and re-assembly of MAGs is outlined in Figure S1. The nitrifier MAGs were subsequently phylogenetically placed in the context of reference *Nitrospira* (Table S4) and *Nitrosomonas* (Table S6) genomes (supplementary text).

### 2.6 Diversity and micro-diversity analysis of nitrifiers

FastANI v1.3 (69) was used to calculate the pairwise average nucleotide identity (ANI) between MAGs within each functional group (i.e., AOB, *Nitrospira-NOB*, and *Nitrospira*-comammox). ANI represents the mean nucleotide identity of the orthologous genes shared between two genomes offering a robust resolution between similar or identical species (i.e., ~80-100% ANI) (69). Since ANI values lower than 80% are not reported, we also calculated the average amino acid identity (AAI) to estimate similarity between two genomes at the amino acid level using compareM (70). To determine intra-population diversity (i.e., microdiversity), we calculated the average nucleotide identity from reads (ANIr) (32) with 90% read identity threshold, as recommended for intra-population comparisons. inStrain v1.3.9 (71) was used to determine the nucleotide diversity for each MAG in each sample and their population average nucleotide identity (popANI) between the samples where they were detected. Nucleotide diversity is a measurement of genetic (micro)diversity at every position along the genome using mapped reads, while popANI is a unique ANI calculation performed by inStrain that considers both major and minor alleles. This is different from the traditional ANI (called consensus ANI [conANI] in inStrain), which only considers major alleles to call (or not) a substitution. Details on parameters and procedures associated with the implementation of ANIr and inStrain are presented in the supplementary text.

### 2.7 Statistical Analyses

Statistical tests were performed using R v4.0.4. All correlations were performed using a linear regression model. The Spearman’s rank correlation coefficients between MAG abundances were performed with the ‘rcorr’ function of the R package Hmisc v4.5.0. Pair-wise significances were calculated with the Kruskal-Wallis test, a nonparametric approach to the one-ANOVA. Principal coordinates analysis (PCoA) with the weighted and unweighted UniFrac distance metrics was used to compare the community composition among systems.

## 3.0 Results

### 3.1 Community composition in nitrogen removal systems

Three nitrification-denitrification systems with high absolute abundance of comammox bacteria were selected for this time-series study on the basis of our previous findings (31) since high sequencing coverage is critical for evaluation of sub-population level diversity. Specifically, over the sampling period (June 2017 to June 2018) comammox bacteria constituted approximately 0.77 (± 0.32), 4.7 (± 4.21), and 0.45 (± 0.24) % of total bacterial 16S rRNA gene abundances at GRE, JAMMSM, and NEU, respectively.

The PacBio full-length 16S rRNA gene sequencing resulted in 2,317,019 total reads (Table S1a) with 7501, 5783, and 8260 ASVs at GRE, JAMMSM and NEU, respectively. Each ASV set was rarefied to the sample with the smallest library size per system resulting in 7040, 4349, and 5749 ASVs from GRE, JAMMSM, and NEU, respectively, and a total of 16651 unique ASVs across all three systems. ASVs with a total of 100 reads in each system were clustered into OTUs at 97% identity, resulting in 846 OTUs (Figure 1A; Table S7). The most abundant OTUs at GRE and NEU were from the classes *Gammaproteobacteria* and *Alphaproteobacteria* and the phylum *Bacteroidota* while the class *Nitrospiria* had the highest relative abundance in JAMMSM. PCoA using weighted (WUF) and unweighted UniFrac (UUF) distance metrics (Figure 1B) demonstrated that samples clustered by system and the community structure between GRE and NEU (WUF: 0.18±0.03; UUF: 0.46±0.03) were significantly more similar than GRE and JAMMSM (WUF: 0.28±0.02; UUF: 0.56±0.05) and NEU and JAMMSM (WUF: 0.29±0.03; UUF: 0.52±0.04) (p<0.05; Figure S2A and B). The greater similarity between GRE and NEU is likely due to the fact that these were suspended phase communities as compared to attached phase (i.e., biofilm samples) communities collected from the IFAS system at JAMMSM.

**Figure 1:**
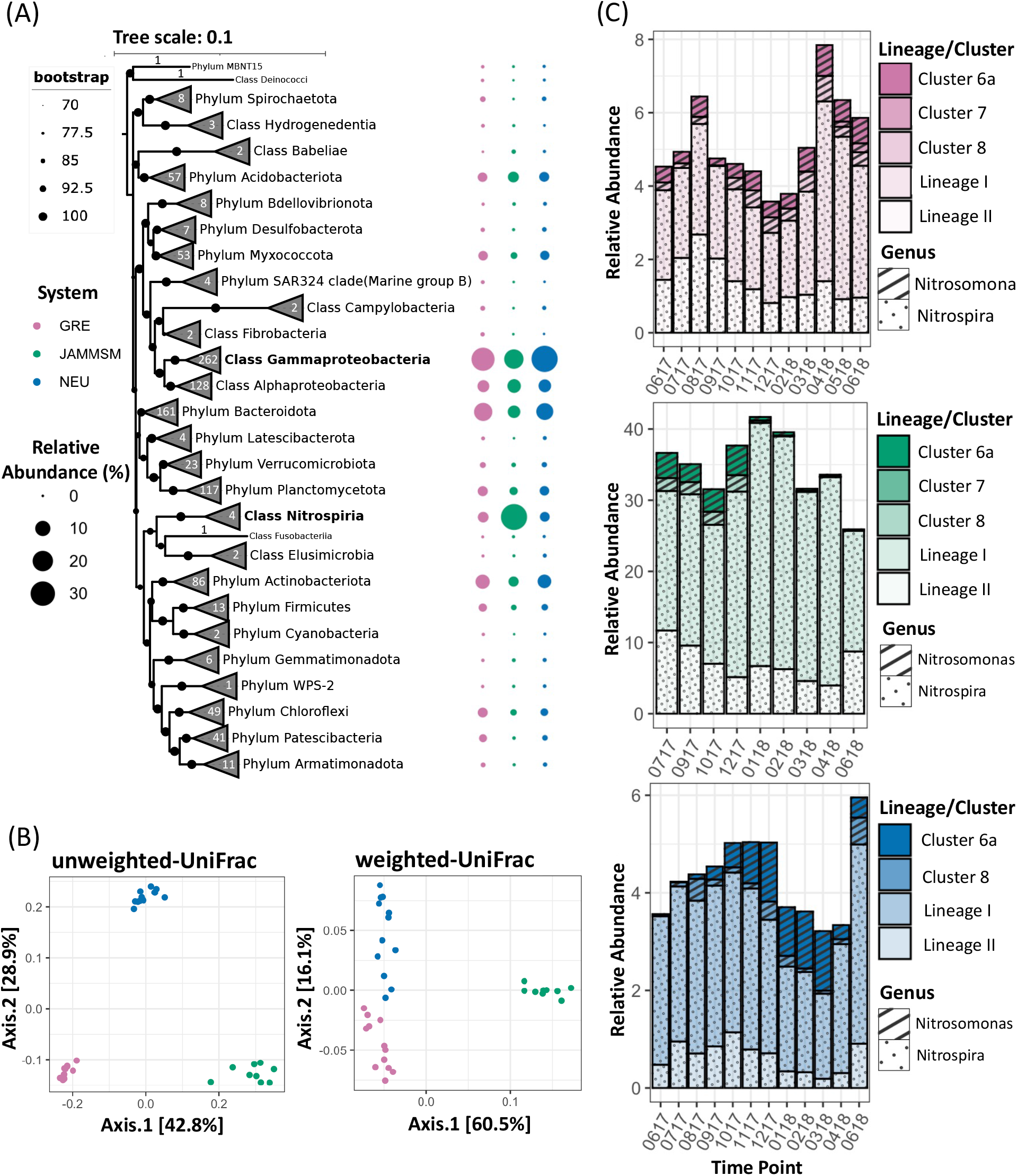
Community composition of the nitrogen removal systems. (A) Phylogenetic tree constructed using full-length 16S rRNA gene sequences of the representative ASV in each OTU (98.7% sequence similarity cluster), with their corresponding relative abundances shown at the right. Branches are collapsed at the phylum/class level and corresponding number of OTUs within each phylum/class across all three systems are indicated. Black circles on branches designate bootstrap support. Bubbles represent cumulative relative abundances of all OTUs within each phylum/class per system. (B) Principal Coordinate Analysis of community composition with unweighted and weighted UniFrac distance metrics using OTU data. Points represent the samples. (C) Cumulative relative abundances of OTUs corresponding to *Nitrospira* Lineages and *Nitrosomonas* clusters per system at each time point analyzed. Colors correspond to each nitrogen removal system with (GRE, pink; JAMMSM, green; NEU, blue).

*Nitrosomonas-* (31, 25, and 14 ASVs at GRE, JAMMSM, and NEU, respectively) and *Nitrospira-like* bacteria (37, 36, and 29 ASVs at GRE, JAMMSM, and NEU, respectively) were the only nitrifiers present in the systems. *Nitrospira* lineage I ASVs were the most abundant group across all systems with average relative abundances of 2.91, 25.13, and 2.88%, followed by *Nitrospira*lineage II with average relative abundances 1.41, 7.07, and 0.64% in GRE, JAMMSM, and NEU, respectively (Figure 1C). *N. oligotropha* (Cluster 6a) and *N. communis* (Cluster 8) -like ASV’s were present in all systems with relative abundances of 0.5 and 0.33, 1.68 and 0.93, and 0.58 and 0.19% in GRE, JAMMSM, and NEU, respectively. *N. europaea/mobilis* (Cluster 7)-like ASVs were also detected in GRE and JAMMSM at very low abundances compared to the other two *Nitrosomonas* linages (average ~0.032 and 0.002%, respectively).

### 3.2 16S rRNA gene sequence-based diversity and microdiversity of nitrifiers

The *Nitrospira-like* ASVs belonged to lineage I (20, 24, and 22 ASVs at GRE, JAMMSM, and NEU, respectively) or lineage II (17, 12, and 7 at GRE, JAMMSM, and NEU, respectively) (Figure S3A), while *Nitrosomonas-like* ASVs were associated with *N. oligotropha* (23, 21, and 10 at GRE, JAMMSM, and NEU, respectively), *N. europaea/mobilis* (5 and 1 at GRE and JAMMSM, respectively), and *N. communis* (3, 3, and 4 at GRE, JAMMSM, and NEU, respectively) lineages (Figure S3B). The *Nitrospira-* and *Nitrosomonas-like* ASVs clustered into 3, 3 and 2, and 21, 14 and 10 OTUs for GRE, JAMMSM and NEU, respectively. The Shannon diversity (i.e., population diversity) of *Nitrosomonas-*like OTUs was significantly higher than *Nitrospira*-like OTUs, while the effective microdiversity of *Nitrospira-like* OTUs was significantly higher than *Nitrosomonas-like* OTUs in all systems (p<0.05; Figure 2A,). *Nitrospira* lineage I OTUs consistently showed the highest microdiversity (Figure 2B) while two *Nitrospira* lineage II OTUs from GRE and JAMMSM had a lower effective microdiversity than most nitrifier OTUs. These results suggest high population level diversity and low microdiversity for *Nitrosomonas*-like bacteria, while the opposite was observed for *Nitrospira-like* bacteria in Lineage I (more ASVs per OTU). The effective microdiversity of *Nitrospira-like* OTUs was positively correlated with their relative abundance (Figure S4A). However, this was neither the case for *Nitrosomonas-like* OTUs (Figure S4B) and the overall bacterial community (Figure S4C). Although the relative abundances of some other highly microdiverse OTUs were positively correlated with effective microdiversity (e.g., *Hyphomicrobium)*, this correlation was not statistically significant for most (e.g., Figure S5). This suggests that the increase in effective microdiversity of *Nitrospira-*like OTUs with increasing relative abundance is more likely to be an ecological phenomenon rather than a sampling artifact.

**Figure 2:**
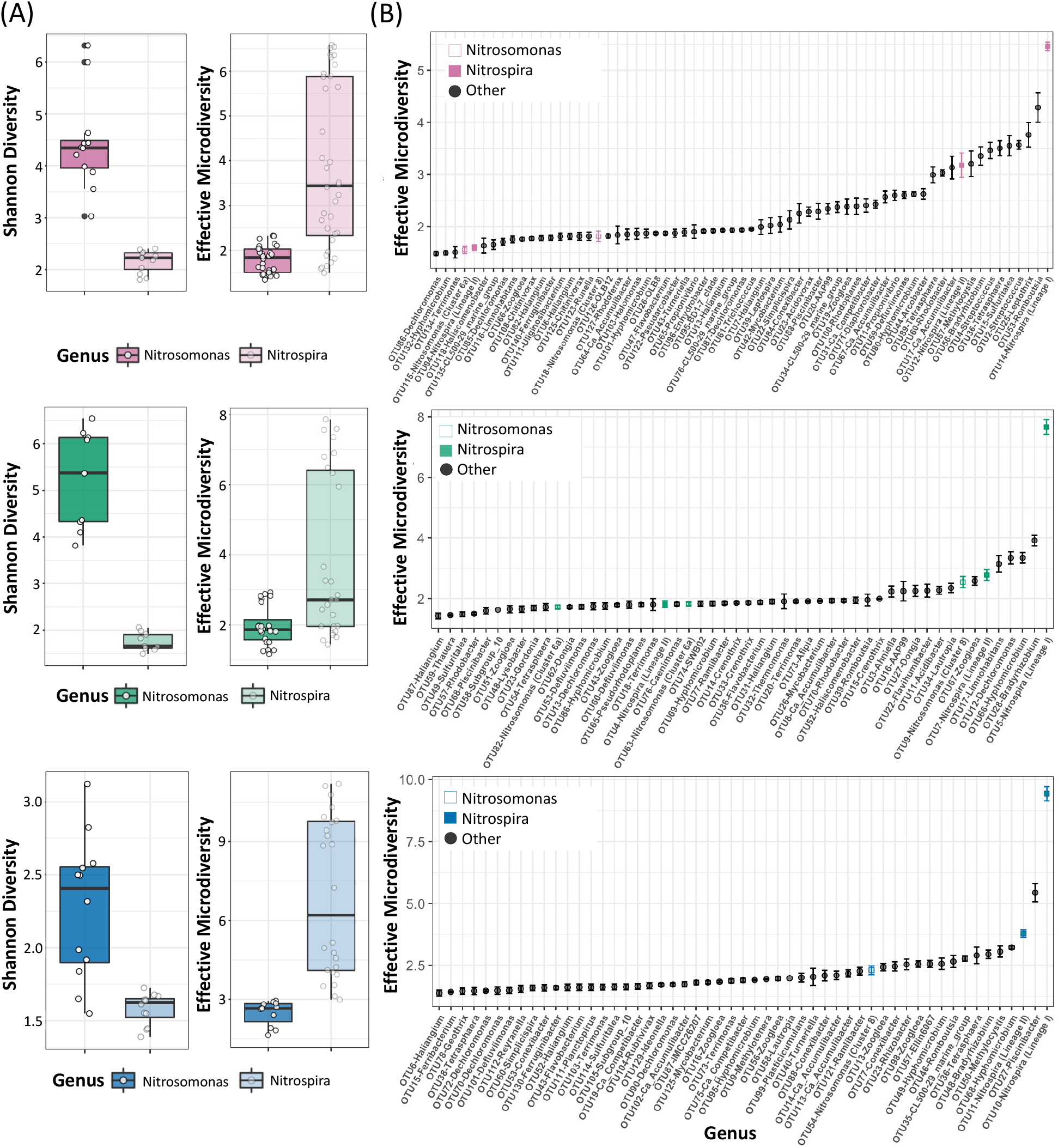
Diversity and microdiversity based on ASV analyses. (A) Boxplot of Shannon diversity and effective microdiversity of *Nitrospira* (lighter shade) and *Nitrosomonas* (darker shade) at each system. (B) Average effective microdiversity of major OTUs per system. *Nitrospira* and *Nitrosomonas* OTUs are colored by system (GRE, pink; JAMMSM, green; NEU, blue), represented by closed and open symbols, respectively.

### 3.3 Improving the recovery of metagenome assembled genomes of nitrifying bacteria

A total of 43 dereplicated nitrifiers MAGS were recovered from short-read assemblies. However, several MAGs lacked key genes associated with nitrification (i.e., *amoA*, *amoB*, *amoC*, *hao*, *nxrA*, *nxrB*). For example, only 2 of the 7 *Nitrosomonas* MAGs (from JAMMSM and NEU) contained the complete *amoCAB* operon and only one had the *hao* gene. To improve MAG quality, we incorporated long reads from Nanopore sequencing and a hybrid assembly using OPERA-MS (37). The hybrid approach increased the contiguity of the assembly and resulted in 30 nitrifier MAGs, with nitrification genes present in 19 MAGs (Table S8). It resulted in significant improvement of MAGs retrieved from JAMMSM but not for GRE or NEU. Thus, we pursued taxonomy aware re-assembly (72) of the 44 non-redundant nitrifer MAGs after dereplicating bins from short-read and hybrid assembly approach. The resultant 36 nitrifier MAGs had completeness greater than 70% and redundancy lower than 10% (Table S8), with nitrification genes present in 24 MAGs. Although the number of retrieved nitrifier MAGs was lower after the reassembly process, there was significant improvement in quality, especially for comammox and AOB genomes. For example, both analyses (before and after reassembly) resulted in 4 *Nitrospira-comammox* MAGs. However, 2 of the 4 comammox MAGs recovered after reassembly contained the entire genetic repertoire required for nitrification, and the other two lack only one gene. In contrast, all pre-reassembly comammox MAGs lacked at least one nitrification gene. These improvements were also observed for AOB including significant improvement in MAG quality (i.e., higher completeness, lower redundancy, and less fragmentation) (Table S8). These improvements resulted from longer contigs created during the reassembly process compared with previous co-assemblies. In contrast to comammox and AOB, despite becoming less fragmented post-reassembly, fewer *Nitrospira*-NOB MAGs contained all genes associated with nitrite oxidation.

### 3.4 Phylogenomic placement and abundance of nitrifier MAGs

The 11 canonical NOB and 4 comammox were associated with *Nitrospira* lineage I (8 *Nitrospira*-NOB) and *Nitrospira* lineage II (3 *Nitrospira*-NOB and 4 *Nitrospira*-comammox), with all comammox MAGs closely related with *Ca*. Nitrospira nitrosa (Figure 3A) and belonged to *Nitrospira-comammox* clade A.1 based on *hao* gene phylogenetic inference (Figure S6). This is in agreement with a previous study that suggests comammox clade A. 1 usually co-occur with the *Nitrospira* lineage I populations in wastewater treatment plants (WWTPs) while clade A.2 and most clade B are typically found in drinking water treatment plants (DWTPs) (21). While non-WWTP settings (e.g., soils, sediments, lakes, rivers, DWTPs) harbor diverse comammox bacteria (9,18,21,73,74), our findings suggest very low diversity of comammox bacteria in wastewater systems which is consistent with other wastewater studies (12,15,17,22,75–77). All AOB MAGs (n=20) were associated with the *N. oligotropha* (n=14) and *N. communis* (n=3) lineages (Figure 3B). Although several members of the *N*. *europaea/mobilis* lineage have been detected in sewage treatment plants (78), only three AOB MAGs related to this lineage were recovered from JAMMSM at very low abundances and less than 50% genome coverage. Thus, these MAGs were excluded from subsequent analyses. The three *N. communis* cluster MAGs (GRE AOB 1, JAMMSM AOB 4, NEU AOB 3) were nearly 100% identical but shared less than 75% ANI with all reference genomes. This lineage can be divided into the *N. communis* and the *N. nitrosa* clusters (79), with *N. communis* species urease negative and detected primarily in agricultural soils, and isolates of *N. nitrosa* urease positive, preferring aquatic habitats and often found in wastewater treatment plants (80). Our three MAGs were urease positive but still distant to *N. nitrosa* in the phylogenomic analysis (Figure 3B), suggesting the presence of a novel phylogenetic lineage. Lastly, most species of the *N. oligotropha* lineage have been recovered from oligotrophic freshwaters and almost all are urease positive (78,80). Accordingly, 12 of the 14 AOB MAGs placed within this *Nitrosomonas* lineage were urease positive.

**Figure 3:**
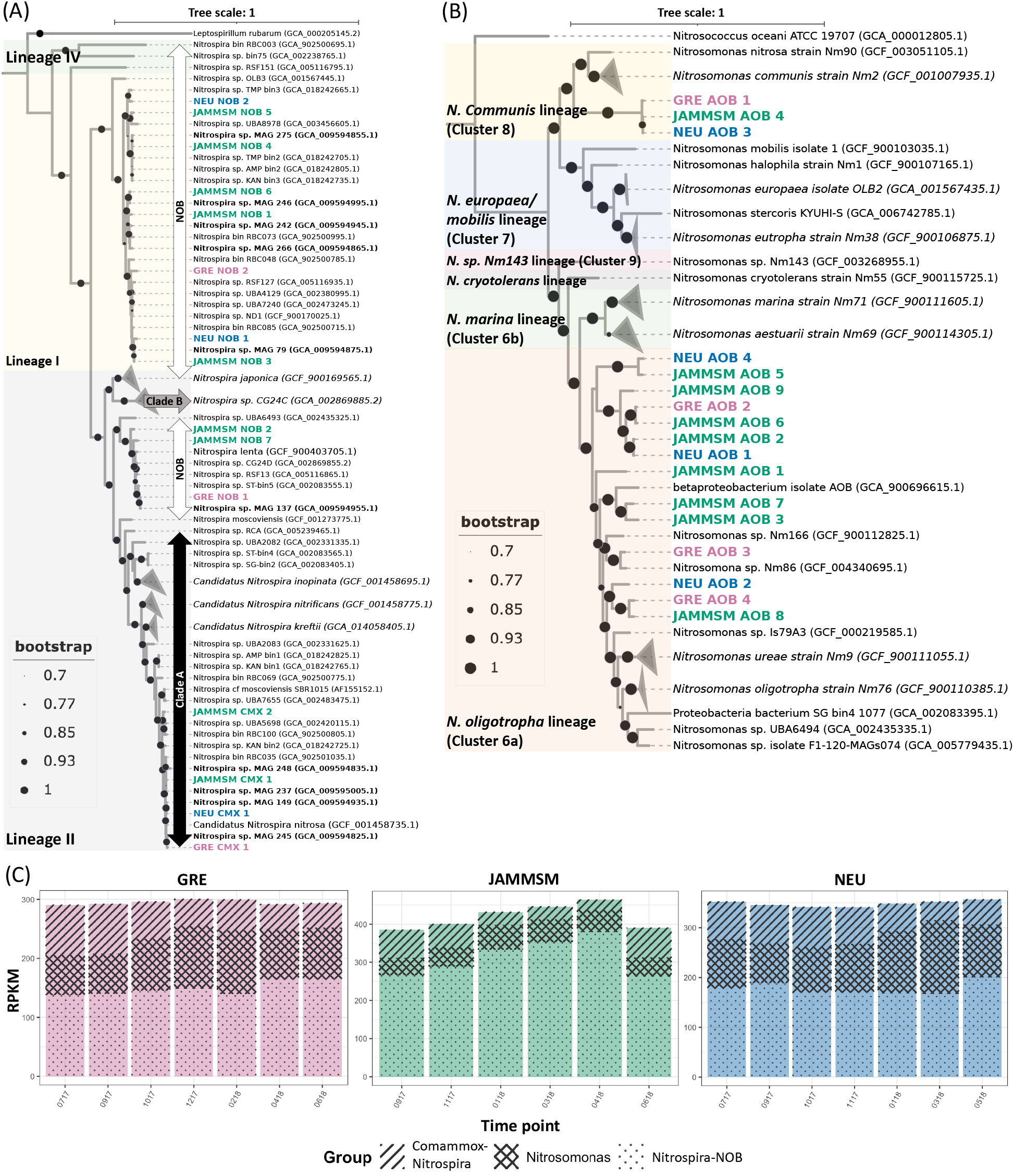
Phylogenetic placement of (A) *Nitrospira* and (B) *Nitrosomonas* MAGs (GRE, pink; JAMMSM, green and NEU, blue) with 78 and 48 reference genomes (black), respectively. Comammox clade A (black arrow), clade B (grey arrow) and canonical NOB (white arrow) are indicated by arrows and phylogenetic lineages by colored boxes. Branches with reference genomes that did not include MAGs from this study were collapsed and labeled with a representative reference genome. All reference genomes used for the reconstruction of the trees are listed in Tables S4 and S6. (C) Cumulative abundances (RPKM) of *Nitrospira*-comammox, *Nitrospira-NOB* and *Nitrosomonas* MAGs at each time point per system.

*Nitrospira*-NOB was the most abundant functional group in all three systems for the duration of the study with higher abundances in JAMMSM (314.5 ±70.6 RPKM) as compared to NEU (178.2 ±12.5 RPKM) and GRE (149.3 ±11.7 RPKM). *Nitrosomonas-like* MAGs were more abundant in NEU (105.2 ±23 RPKM) and GRE (86 ±15.8 RPKM) as compared to JAMMSM (53.2 ±27.4). This suggests that *Nitrospira*-like bacteria may prefer biofilm-based growth over suspended phase (81). *Nitrospira*-comammox relative abundances were very similar in all systems, ranging from 55.5 to 64.9 RPKM in JAMMSM and NEU, respectively. The abundances of nitrifying MAGs were weakly correlated with each other, with only 37 of 195 correlations significant (p-value <0.05). Most of the significant correlations between AOB and *Nitrospira*-NOB were positive (Figure S7), likely arising as consequence of metabolic interactions of these groups within the nitrification process. Comammox bacteria had very few correlations with other nitrifiers, with primarily negative correlations with MAGs from *N. oligotropha*-like bacteria (cluster 6a).

### 3.5 Population-level diversity of nitrifying bacteria within and between systems

Post dereplication at 95% ANI (69,82), pairwise ANI and AAI were calculated between all MAGs within the same functional group (i.e., AOB, *Nitrospira-NOB*, and *Nitrospira-comammox)*(Figure 4A-B, Tables S9a-b). Of the 136 pairwise comparisons between the 17 AOB MAGs, only 18 (13.2%) had ANI values higher than 80% suggesting high population-level diversity within and between nitrogen removal systems for the *Nitrosomonas*-like AOB (Figure 4A). The pairwise AAI for *Nitrospira*-NOB MAGs showed a bimodal distribution (Figure 4B) reflective of the two lineages detected in each system (i.e., Lineage I and II). Pairwise AAIs for *Nitrospira-NOB* MAGs of the same lineage within or across systems were typically >85% (Table S9b). These results suggest that while multiple AOB and *Nitrospira*-NOB coexist in each system, AOB were substantially more diverse than *Nitrospira-NOB*. In contrast, three of the four comammox MAGs (one from each system) share ANI values between 95-97% and are closely related with *Ca.*Nitrospira nitrosa (31). Although a comammox MAG from JAMMSM (JAMMSM CMX 2) has lower than 95% with the other three MAGs, it still exhibited greater than 85% AAI (Figure 4B) with them and also falls within the *Ca*. Nitrospira nitrosa cluster (Figure 3A)(83). Pairwise AAI comparisons (which are highly correlated with ANI – Figure S8) indicated that *Nitrospira-*comammox MAGs have significantly higher AAI compared with AOB and NOB, while AOB were the most diverse functional group at the population level (p <0.05; Figure 4B). Moreover, similar to ASV analysis, *Nitrospira*-NOB MAGs were associated with one of two clusters (*Nitrospira* lineage I or II), with highly similar populations within each cluster (Figure 4B) being detected across systems. These findings further confirm that in contrast to AOB that demonstrate high population diversity, comammox bacteria show very low population diversity and, in fact, may all be associated with the same population across multiple wastewater systems

**Figure 4:**
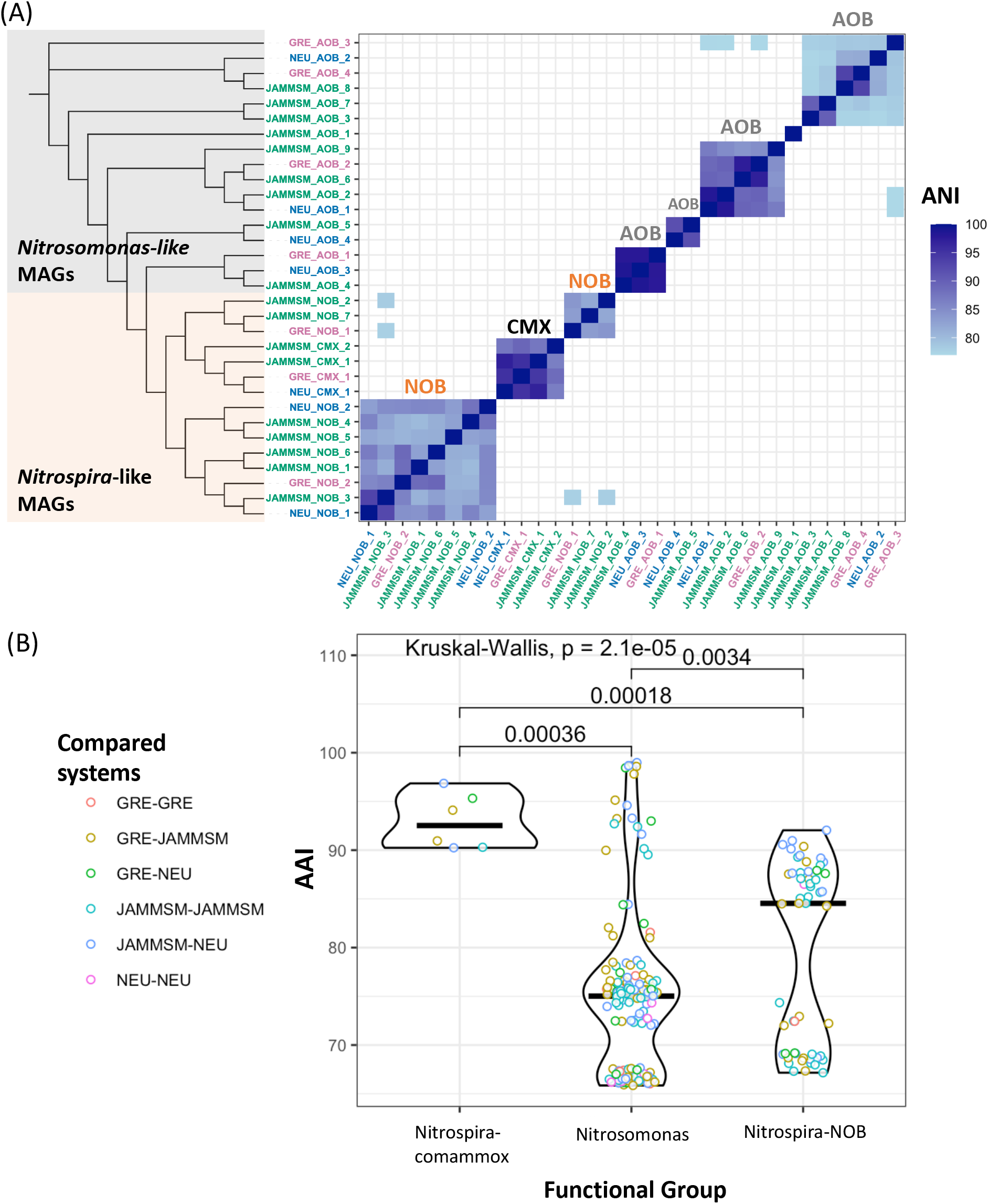
(A) ANI values calculated between *Nitrosomonas* (AOB), *Nitrospira-NOB* and *Nitrospira-comammox* (CMX) MAGs along with the phylogenetic placement of the MAGs. MAG labels are colored by system (GRE, pink; JAMMSM, green and NEU, blue). (B) AAI values per functional group. Black lines represent the average, points represent the AAI value between two MAGs and colors decode the systems that are compared. Pairwise statistical comparisons were performed with the Kruskal-Wallis test.

### 3.6. Microdiversity of nitrifying bacteria

The ANIr distributions for comammox and AOB MAGs were not significantly different from each other, and significantly higher than for *Nitrospira-NOB* MAGs (Figure 5A). While *Nitrospira-*NOB overall demonstrated lower ANIr indicating higher microdiversity compared to AOB and comammox bacteria, there was some variability within each system (Figure S9). These observed trends were similar between ANIr and inStrain estimated nucleotide diversity (Figure 5B), with AOB showing on average lowest microdiversity and *Nitrospira-NOB* the highest. Specifically, the average nucleotide diversity of *Nitrosomonas*-like AOB was significantly lower than of both *Nitrospira* groups, i.e., NOB and comammox (p <0.05), while with ANIr only *Nitrospira-NOB*were significantly more microdiverse than the two other functional groups. This difference arises from the fact that the two measures of intra-population diversity (i.e., ANIr and nucleotide diversity) are calculated differently. For instance, ANIr considers only the major alleles in the consensus sequence to call a substitution, while nucleotide diversity is calculated using base pair frequencies at each position. Nonetheless, both approaches indicate that *Nitrospira*-NOB exhibit significantly higher microdiversity as compared to *Nitrosomonas*-like AOB and *Nitrospira*-comammox bacteria. These results are consistent with the 16S rRNA gene analysis where *Nitrospira* lineage I and most lineage II OTUs had significantly higher effective microdiversity than the *Nitrosomonas*-like OTUs and the remaining *Nitrospira* lineage II OTUs. Based on these results, we speculate that the *Nitrospira* lineage II OTUs with low effective microdiversity likely correspond to comammox bacteria (Figure 2B).

**Figure 5:**
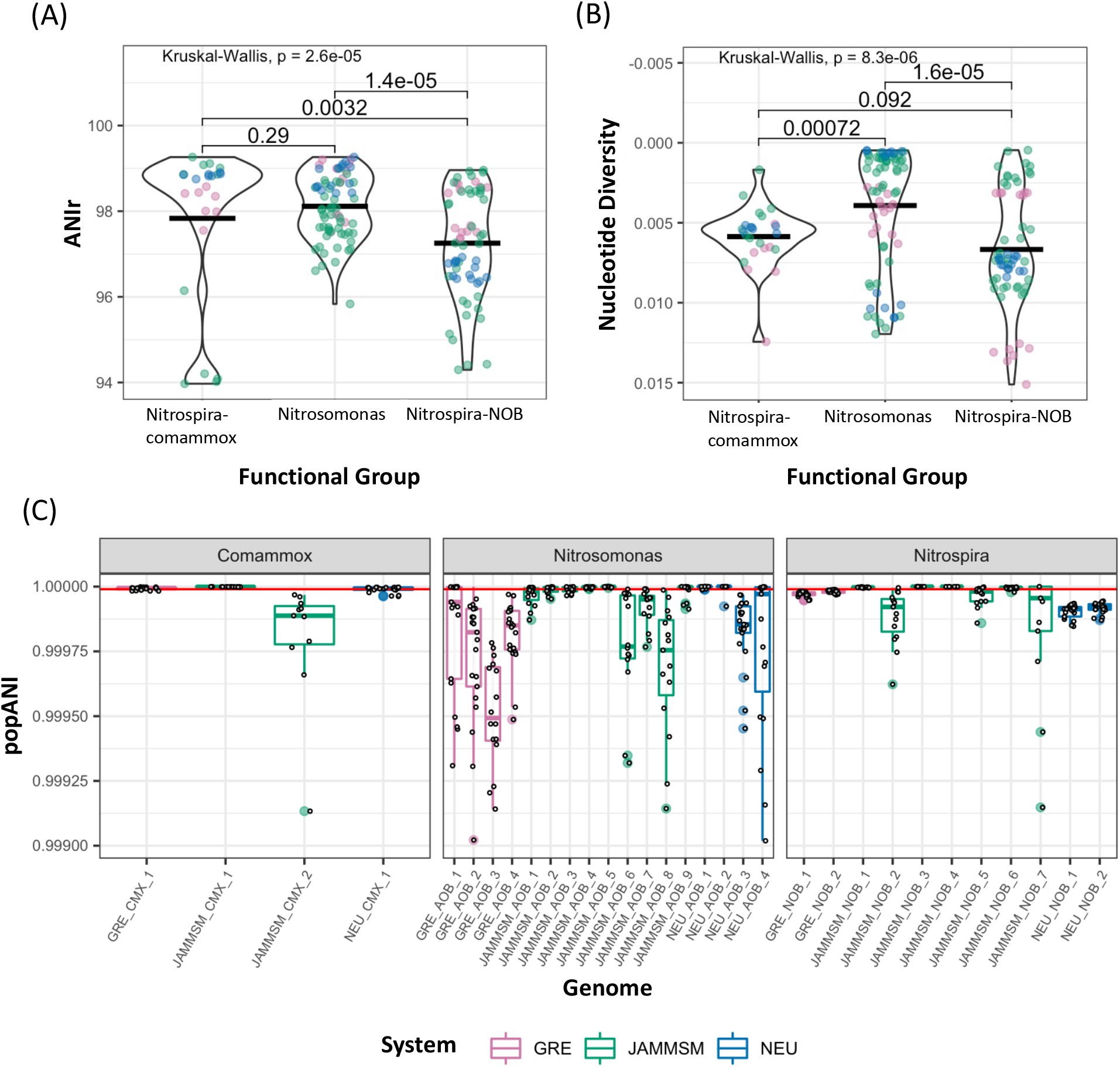
(A) ANIr and (B) nucleotide diversity values from each functional group. Points represent the average value of each MAG and are colored according to the systems (GRE, pink; JAMMSM, green and NEU, blue) from which the MAGs were recovered. (C) popANI values from each MAG. MAGs are colored by system and points represent the popANI value of the MAG between two samples.

Finally, popANI, a microdiversity-aware ANI calculation, was used to discriminate between strains across samples using the recommended popANI threshold of 99.999% (Olm et al., 2021). All popANI values from GRE_CMX_1, JAMMSM_CMX_1, and NEU_CMX_1 were above or very close to the 99.999% threshold, suggesting low strain-level diversity of comammox bacteria (Figure 5C). Only one comammox genome retrieved from JAMMSM (JAMMSM_CMX_2) had popANI values lower than the recommended threshold. However, the average relative abundance of this comammox (RPKM = 7 ±2) was particularly low as compared to the more abundant comammox in JAMMSM (RPKM = 48 ±21) and those retrieved from GRE (RPKM = 60 ±19) and NEU (RPKM = 65 ±17). Therefore, the principal or sole comammox bacteria in each system is not only a single population but likely also a specific strain. The ANIr for each comammox MAG estimated by mapping reads from a different system (e.g., mapping GRE or NEU reads to MAGs assembled from JAMMSM) were lower than when mapping of reads from the system from where the MAG was obtained (e.g., mapping JAMMSM reads to MAGs assembled from JAMMSM; Figure S10). This suggests that on the one hand the comammox bacterial population within a single system is restricted to a specific *Ca*. Nitrospira nitrosa-like strain, but on the other hand that different systems contain different strains.

## 4 Discussion

### 4.1 Hybrid assembly and re-assembly processes improve the quality of metagenome assembled genomes

Reliance on short reads for genome assembly from complex metagenomes can lead to incomplete and fragmented MAGs especially in systems containing closely related populations (84). Indeed, short read analysis resulted in highly fragmented nitrifier MAGs lacking several nitrification genes. To improve the short read assemblies, we used long reads generated from Nanopore sequencing with a hybrid assembly approach (38). Recent studies have demonstrated the ability of hybrid assemblies to significantly improve the recovery of complex bacterial genomes (85,86) by increasing the contiguity and accuracy of metagenome assembly (87). Combining the two sequencing techniques (i.e., short and long reads) with the hybrid assembler OPERA-MS resulted in modest assembly and binning improvements regarding genome contiguity and completeness. However, the bead beating based DNA extraction protocol may have resulted in shearing of genomic DNA into smaller fragments which is not ideal for long-read sequencing (88) and resulted in lower read lengths. In an attempt to further improve MAG quality, we performed taxonomy aware reassemblies (72) as described in the materials and methods section. Since a hybrid approach resulted in some improvements in MAG quality, we also adopted a hybrid strategy with OPERA-MS for taxonomy aware reassembly. This significantly improved the completeness and contiguity of MAGs and number of annotated nitrifier genes (especially for *Nitrosomonas* and comammox *Nitrospira*) binned within them. While the *Nitrospira*-NOB MAGs also became less fragmented, they showed fewer improvements with respect to the presence of nitrite oxidation genes. This is likely caused by the high levels of microdiversity within *Nitrospira-NOB* (Figures 2 and 5), which likely impacts both the read mapping and de novo assembly process. In the future, this issue might be improved by incorporating more long-read data into the re-assemblies, especially once the accuracy of long-read sequencing technologies further increases.

### 4.2 *Ca*. Nitrospira nitrosa-like bacteria are the primary comammox bacteria in secondary wastewater treatment for nitrogen removal

Structural diversity and functional redundancy are inherently linked to process stability when environmental and process conditions vary (28). Both, metagenomics and 16S rRNA gene sequencing, indicated that *Nitrosomonas* and *Nitrospira*-like bacteria were the only known nitrifiers present in the systems investigated. This is consistent with numerous studies (26,35,89,90) suggesting the specific adaptation of these genera to the wastewater environment. The majority of the *Nitrosomonas-like* AOB MAGs, within and between systems, had AAI values below 85% (Figure 4) indicating that this functional group exhibited high species level diversity. Meanwhile, *Nitrospira*-like MAGs within the same lineage reported AAI values above 85% in most cases. The high population level diversity of the *Nitrosomonas*-like bacteria was also confirmed with full length 16S rRNA gene sequencing (Figure S3B) and corroborates earlier studies on their diversity in wastewater (26,90,91). Although *N. europaea*, *N. oligotropha* and the *Nitrosomonas*-like clusters closely associated with *N. communis*/*nitrosa* are frequently the most abundant groups in wastewater, different *Nitrosomonas* populations dominate different processes and the extent of their diversity is system dependent (26). For example, *N. europaea* are often the major AOB in high ammonia concentration environments (e.g., anammox reactors) due to low ammonia affinity and low DO level adaptation (91). The observed dominance of *N. oligotropha*and *N. communis* in this study are consistent with previous studies of WWTPs with relatively lower ammonia concentrations (26,91,92).

Comammox bacteria exhibited significantly lower population level diversity as compared to strict AOB and NOB (Figure 4). All comammox *Nitrospira* MAGs, independent of the system they were recovered from, belong to *Nitrospira* lineage II and were closely associated with *Ca*. Nitrospira nitrosa. These findings are consistent with several other studies that have found *Ca*. Nitrospira nitrosa-like bacteria as the unique or principal comammox bacteria present in their laboratory and full-scale wastewater treatment systems (11,15,17,20,22,75,76), both using metagenomics and gene targeted analyses. In contrast, other engineered and natural habitats typically harbor multiple co-existing species across the two primary clades (i.e., comammox clade A and B). For example, a study of multiple groundwater-fed sand filters recovered 12 comammox MAGs (5 clade A and 7 clade B) and revealed the co-occurrence of at least five comammox species (10 on average) per system (34). Other studies in soils, sediments, lakes, drinking water, groundwater, and estuaries have also reported the co-existence of multiple comammox species (9,73,93,94). However, this co-existence of comammox populations across clades does not seem to be prevalent in wastewater treatment systems. For instance, a study of 8 WWTPs found 9 of 14 comammox OTUs were associated with the *Ca*. Nitrospira nitrosa cluster and accounted for 94.34% of all comammox *amoA* sequences (17) with different OTUs representing the dominant comammox species in different systems. Moreover, Beach and Noguera (75) reported the dominance of *Ca*. Nitrospira nitrosa in five low DO nitrogen removal bioreactors using qPCR with species-specific primers. These findings suggest that the diversity of comammox bacteria in WWTPs is lower than in other habitats where several comammox species co-exist.

In contrast, Spasov et al. (16) reported the detection of multiple comammox MAGs in samples collected from rotating biological contactors (RBCs) used for tertiary treatment at a municipal WWTP in Ontario, Canada. The difference in population diversity between that study and other WWTP investigations, including this one, may be associated with ammonia availability. Specifically, tertiary treatment systems are typically employed as a polishing step; they are designed and operated to remove low residual amounts of nitrogen. The influent ammonium concentrations reported for the RBC tertiary treatment system range from 0.2 to 16.3 μM (16), while the three systems in this study have influent ammonia concentrations between 2 and 3 mM. The comammox *Nitrospira* MAGs with overall higher abundances in the tertiary plant (RBC001 and RBC083) were less abundant at the beginning of the train (higher ammonium concentrations) than at the end of the train (lower ammonium concertation). In contrast, the abundance of RBC035, the only MAG phylogenetically associated with *Ca*. Nitrospira nitrosa (ANI >95%), decreased in abundance with decreasing ammonium concentrations. The high comammox diversity in the RBCs and other ammonium-limited systems suggest that ammonium concentration and/or flux through the system may be a key factor driving comammox bacteria diversity in nitrification systems. In fact, Palomo et al. (34) reported that the influent ammonium concentration was the key explanatory variable associated with comammox bacterial diversity in 12 groundwater-fed sand filters when higher comammox population diversity was detected in systems receiving lower ammonium concentrations. *Ca*. Nitrospira inopinata and *Ca*. Nitrospira kreftii, two comammox species with known kinetic parameters, have high apparent ammonia affinities (95,96); assuming this is a conserved trait in comammox bacteria, this explains their prevalence and diversity in ammonium-limited systems. However, the kinetic parameters (i.e., affinity constant, growth rate) of other comammox species especially of the *Ca*. Nitrospira nitrosa clade have not yet been reported, and our findings suggest that at least this comammox clade may have substantially different kinetic traits that allow it to thrive in systems with higher ammonium concentrations. Low DO has also been associated with the prevalence of comammox bacteria over canonical nitrifiers in wastewater systems (15,75). However, the high DO concentrations (>2 mg/L) in this and other studies (17,20) suggest DO is not a crucial factor for the proliferation of *Ca*. Nitrospira nitrosa in nitrogen removal systems.

### 4.3 Comammox bacteria not only exhibit low diversity at the species/population level but also at the strain/intra-population level across wastewater systems

The high abundance and microdiversity of *Nitrospira*-NOB have been previously reported in wastewater and other engineered and natural environments (4,33,35). However, it is unclear whether the high microdiversity of *Nitrospira* is associated with high functional diversity or is indicative of the coexistence of functionally identical *Nitrospira* with allelic diversity. Gruber-Dorninger et al. (35) reported variable responses of closely related bacteria within *Nitrospira*lineage I (sequence identities ranging from 95.8 to 99.6%) to different nitrite availabilities. Similarly, ecological niche partitioning was also identified as potential mechanisms for co-existence of three *Nitrospira* lineage I strains that used formate under different conditions: two used formate when incubated with nitrite and ammonia, respectively, while the third used formate efficiently as the sole substrate (35). Although these results shed light on the possible niche partitioning of *Nitrospira* sub-species, this does not shed light on whether the microdiversity in our study represents coexisting *Nitrospira* strains with slight functional differences. Although functional implications of such high microdiversity are not clear, the persistence of high intra-population diversity may suggest it plays an important role in the distribution and success of *Nitrospira* populations in wastewater systems. A few studies in other environments have shown that persistent populations exhibit increased intra-population sequence diversity (7,32). A study in a saltern pond suggested that the ecologically important genes of the major archaeal sequence-discrete population were carried by distinct sub-populations (strains), indicating that the adaptation to different salinity concentrations had led to sub-population differentiation and speciation (niche partitioning) (82). Moreover, a study in eight different temperate bog lakes concluded that high microdiversity is associated with the maintenance of functional microbial communities during changes in environmental conditions (36).

Contrastingly, comammox bacteria not only exhibited an unexpected low population-level diversity within and between systems but also showed significantly lower microdiversity as compared to strict NOB within the genus *Nitrospira*. These results may indicate specific adaptations of comammox bacteria within the *Ca*. Nitrospira nitrosa cluster to the wastewater environment and particularly secondary treatment systems. Palomo et al. (34) observed a negative correlation between the species-level diversity of comammox *Nitrospira* and ammonium concentrations. However, that was not the case for microdiversity, suggesting that different mechanisms may shape inter-versus intra-population diversities, or that the range of ammonium concentrations in the investigated drinking water systems was too narrow to capture any underlying associations. For instance, the comammox nucleotide diversity in our study ranged from 0.002 to 0.008, while this was from ~0.005 to ~0.013 for the MAGs obtained from these drinking water systems, with only three of the twelve comammox MAGs presenting nucleotide diversities lower than 0.008 (34). This may suggest that comammox microdiversity is also associated with ammonium concentrations and/or flux through the system, with the observed low microdiversity in wastewater systems associated with the higher prevailing ammonium concentrations.

Despite the broad detection of comammox bacteria in WWTPs, their role and process relevance are as yet unclear. In contrast to other environments, comammox *Nitrospira* present very low population and intra-population diversity levels in wastewater treatment systems for nitrogen removal. This observed lack of diversity and, consequently, lack of functional redundancy may influence the feasibility of potential design and operational strategies relying primarily on comammox *Nitrospira* for nitrogen removal. Although ammonium concentration and/or availability apparently influences the diversity of comammox bacteria, further studies are necessary to determine other factors driving the success of their clonal community in wastewater. Moreover, studies are required to estimate the activity of comammox bacteria, specifically *Ca*. Nitrospira nitrosa, and to assess their contribution to nitrification in full-scale nitrogen removal systems. It is also important to note that the intra-population diversity cannot be solely studied with shotgun metagenomic methods since high microdiversity can prevent robust assembly of individual genomes (82). Therefore, more accurate techniques are needed to obtain high-quality strain-level genome assemblies for *Nitrospira* and other highly microdiverse bacteria.

## Supporting information

Supplemental Tabel 1

Supplemental Tabel 2

Supplemental Tabel 3

Supplemental Tabel 4

Supplemental Tabel 5

Supplemental Tabel 6

Supplemental Tabel 7

Supplemental Tabel 8

Supplemental Tabel 9

## 4 Acknowledgements

This research was supported by the National Science Foundation (Award number: 1703089) and the Netherlands Organization for Scientific Research (016.Vidi.189.050). The authors also acknowledge the operational personnel for assistance with sampling and process data sharing. The authors also thanks A.M. Eren for assistance with metagenomic co-assemblies.

## 5 Data availability

All raw sequencing data from Illumina and Nanopore platforms and nitrifiers MAGs are available on NCBI under bioproject number PRJNA846349.

