## Supplemental Figures for "Low diversity and microdiversity of comammox bacteria in wastewater systems suggests wastewater-specific adaptation within the *Ca*. Nitrospira nitrosa cluster"

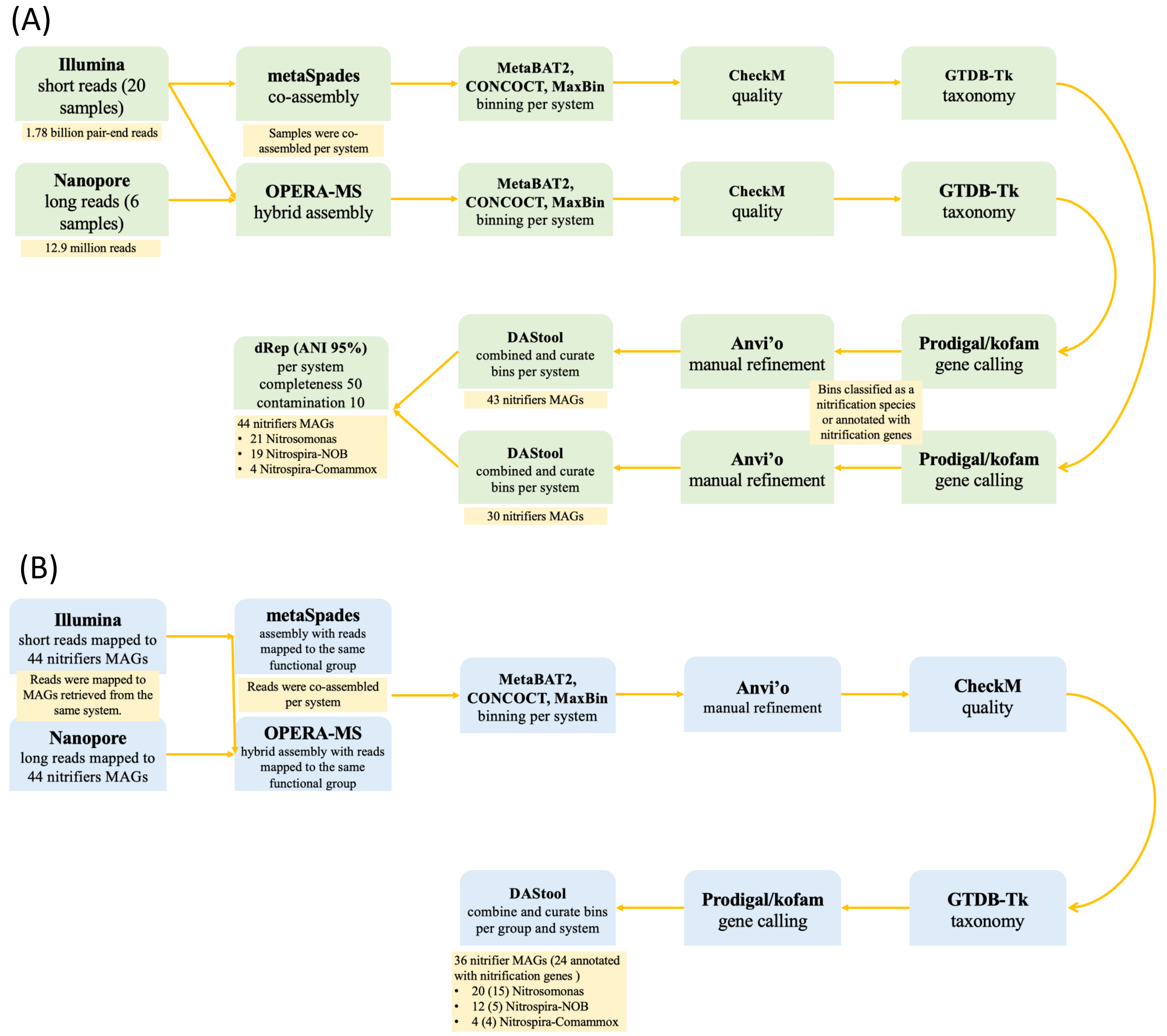


**Figure S1:** MAG generation workflow for de-novo assembly (A) and targeted nitrifier re-assembly (B). Co-assemblies were performed by system and re-assemblies by system and functional group.

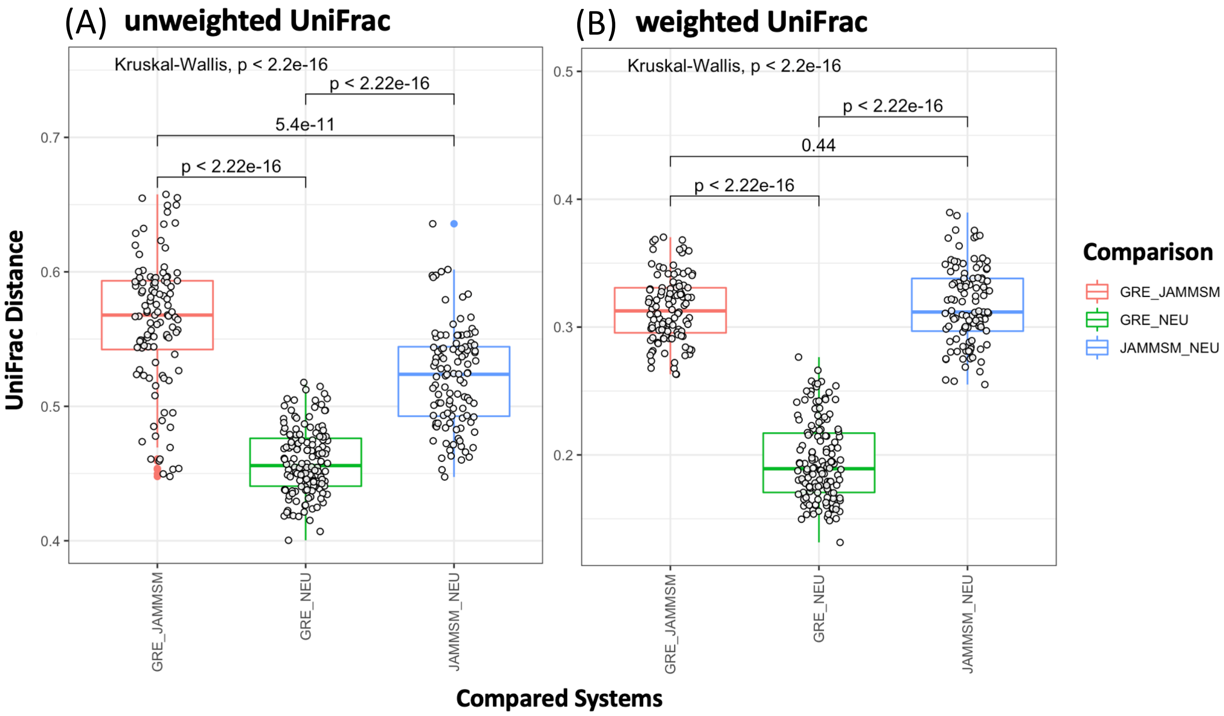


**Figure S2:** Pairwise unweighted (A) and weighted (B) UniFrac distances between systems.

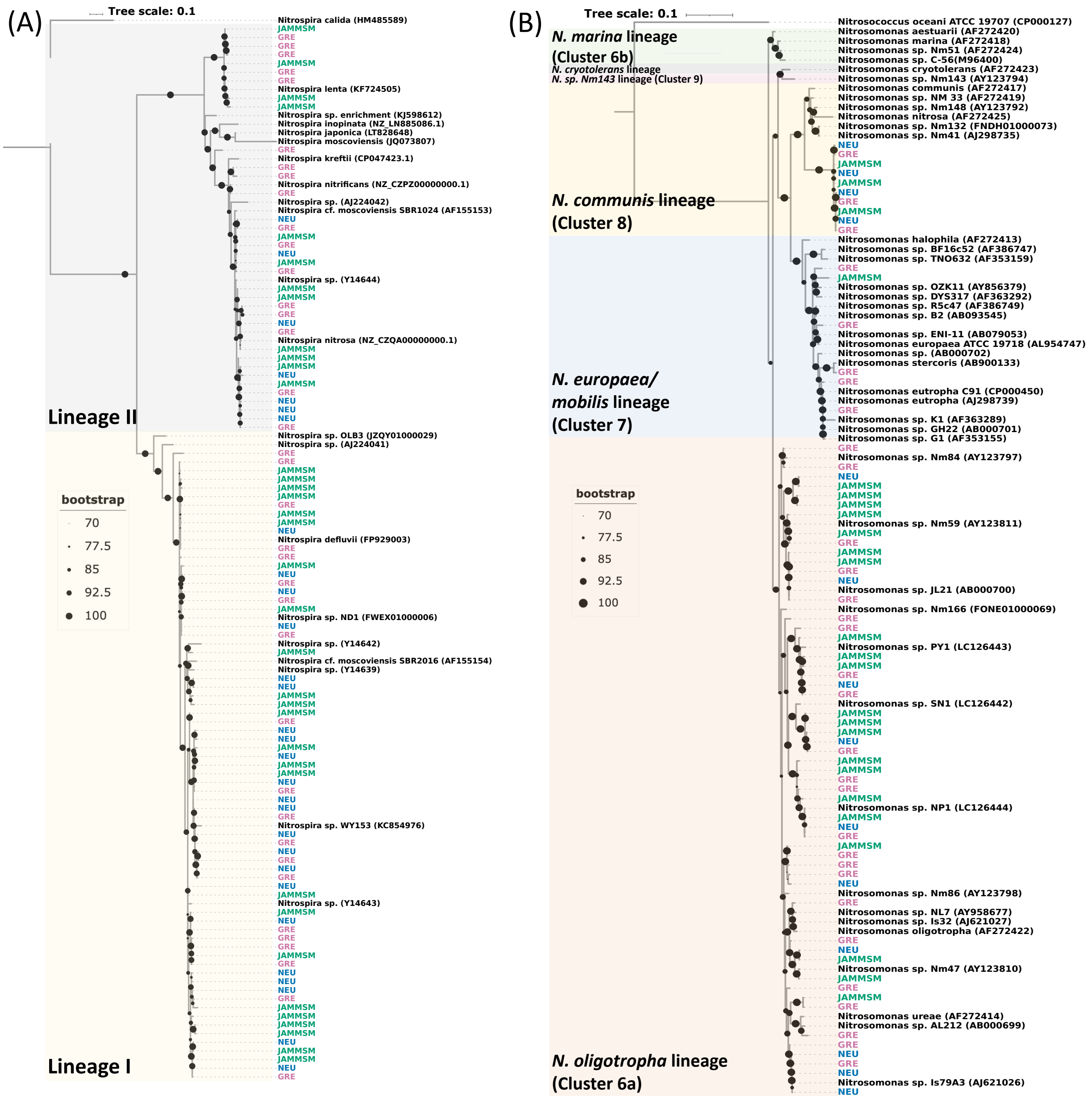


**Figure S3**: Phylogenetic tree of the 16S rRNA gene sequences from (A) *Nitrospira*-like and (B) *Nitrosomonas*-like ASVs along with reference sequences labeled in black. Clades are colored by Lineage/Cluster and ASVs recovered from this study are colored by system (GRE, pink; JAMMSM, green; NEU, blue).

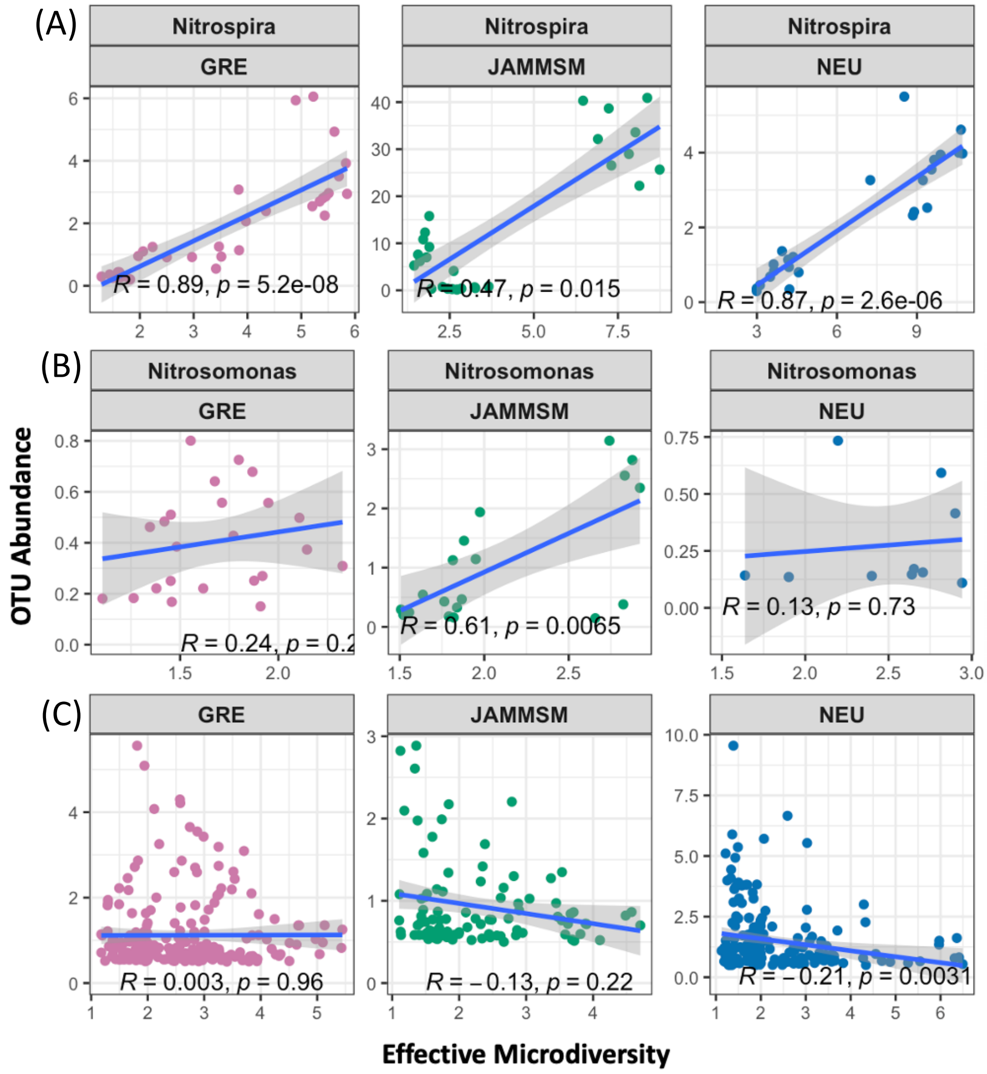


**Figure S4:** Correlation between relative OTU abundance vs. effective microdiversity of (A) *Nitrospira*-like bacteria, (B) *Nitrosomonas*-like bacteria, and (C) all OTUs with relative abundance greater than 0.1% in the bacterial community (except for nitrifiers). Data points represent OTUs with abundances higher than 0.1% and are colored by system.

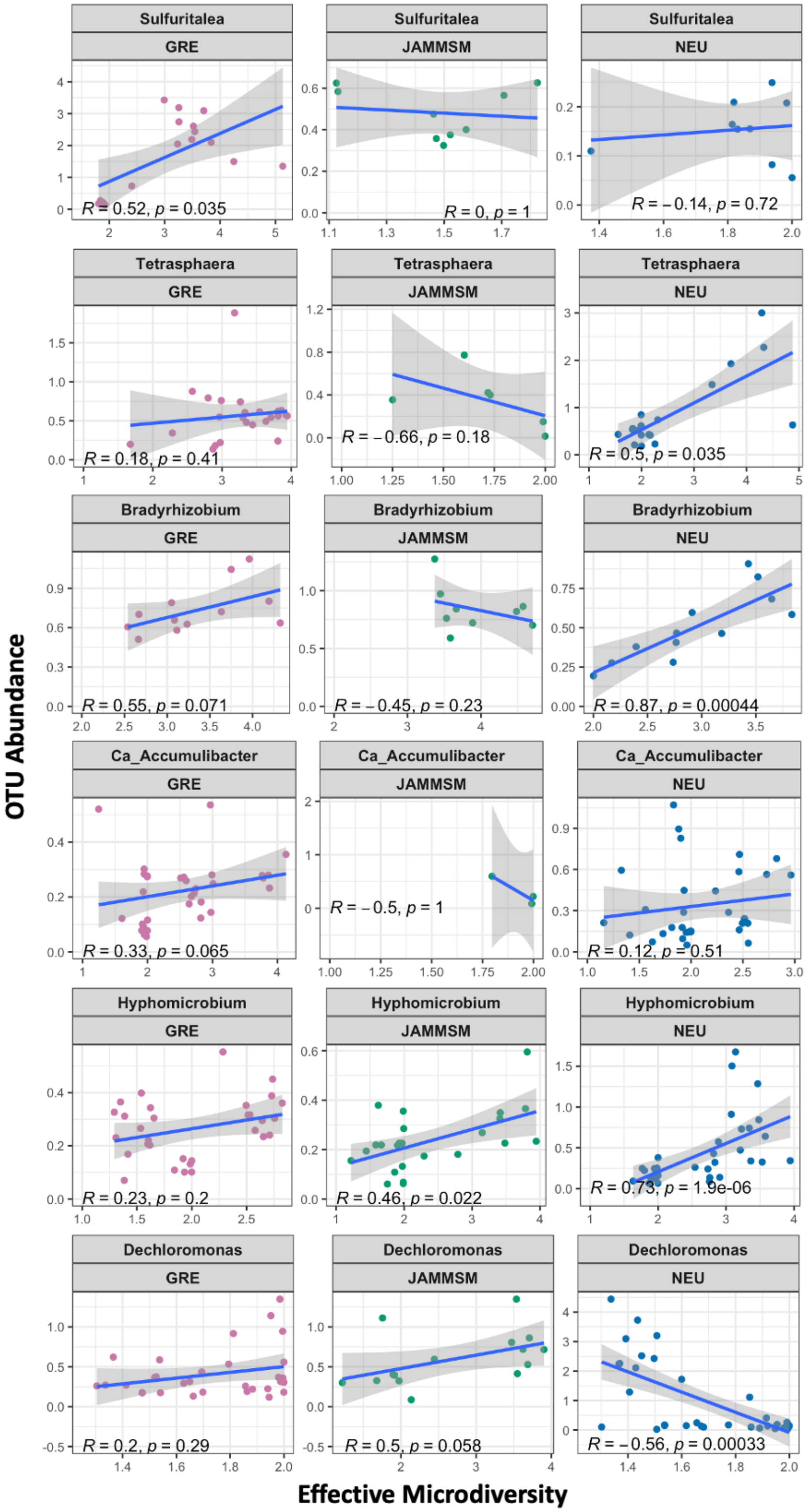


**Figure S5:** OTUs relative abundances vs effective microdiversity of most microdiverse taxa per system. Data points represent OTUs with abundances higher than 0.1% and are colored by system.

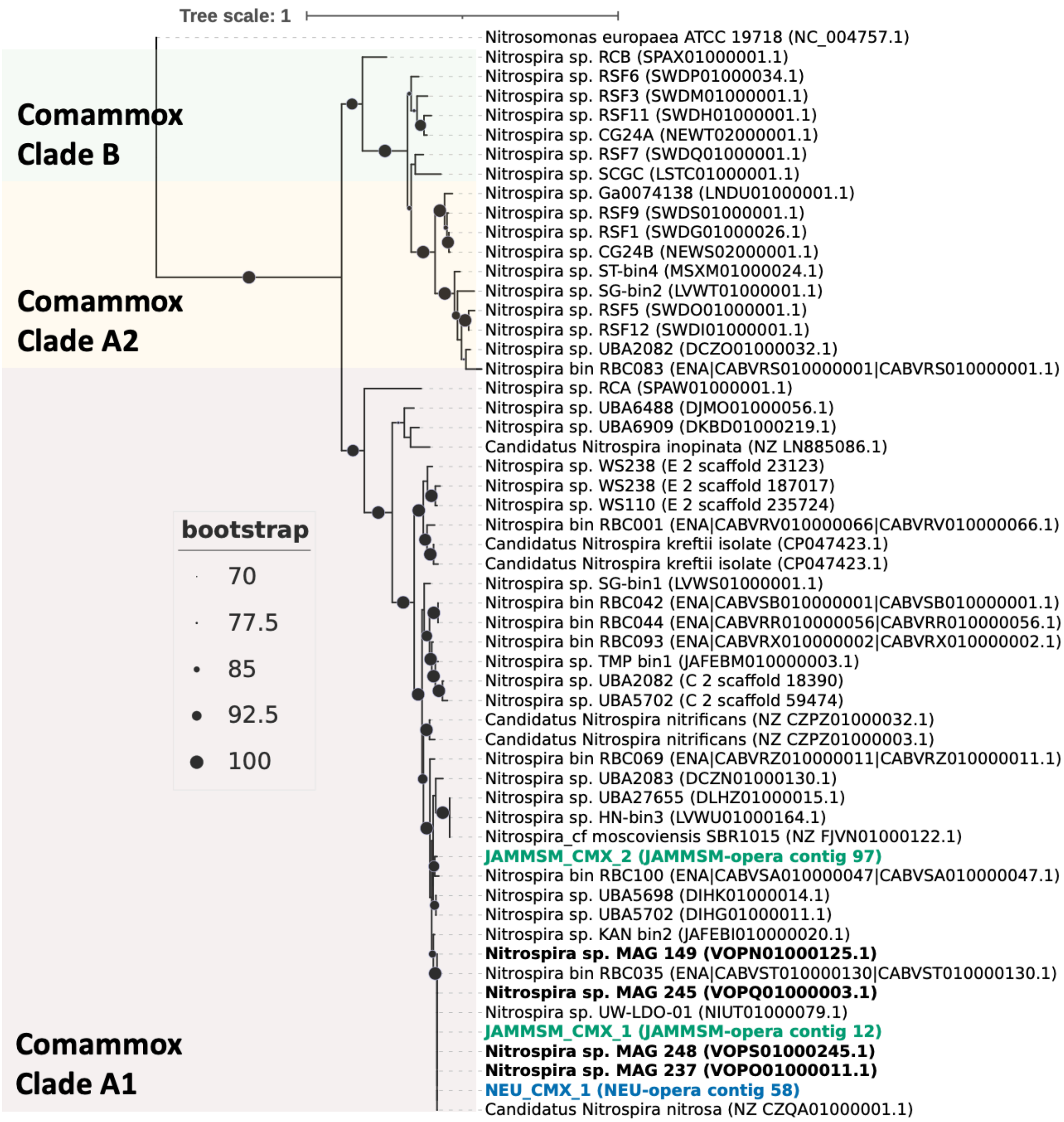


**Figure S6:** Maximum likelihood phylogenetic tree of comammox *Nitrospira* MAG hao genes colored by system (JAMMSM, green; NEU, blue) with 48 comammox *Nitrospira hao* references genomes (black). Black references in bold are sequences for our previous study.

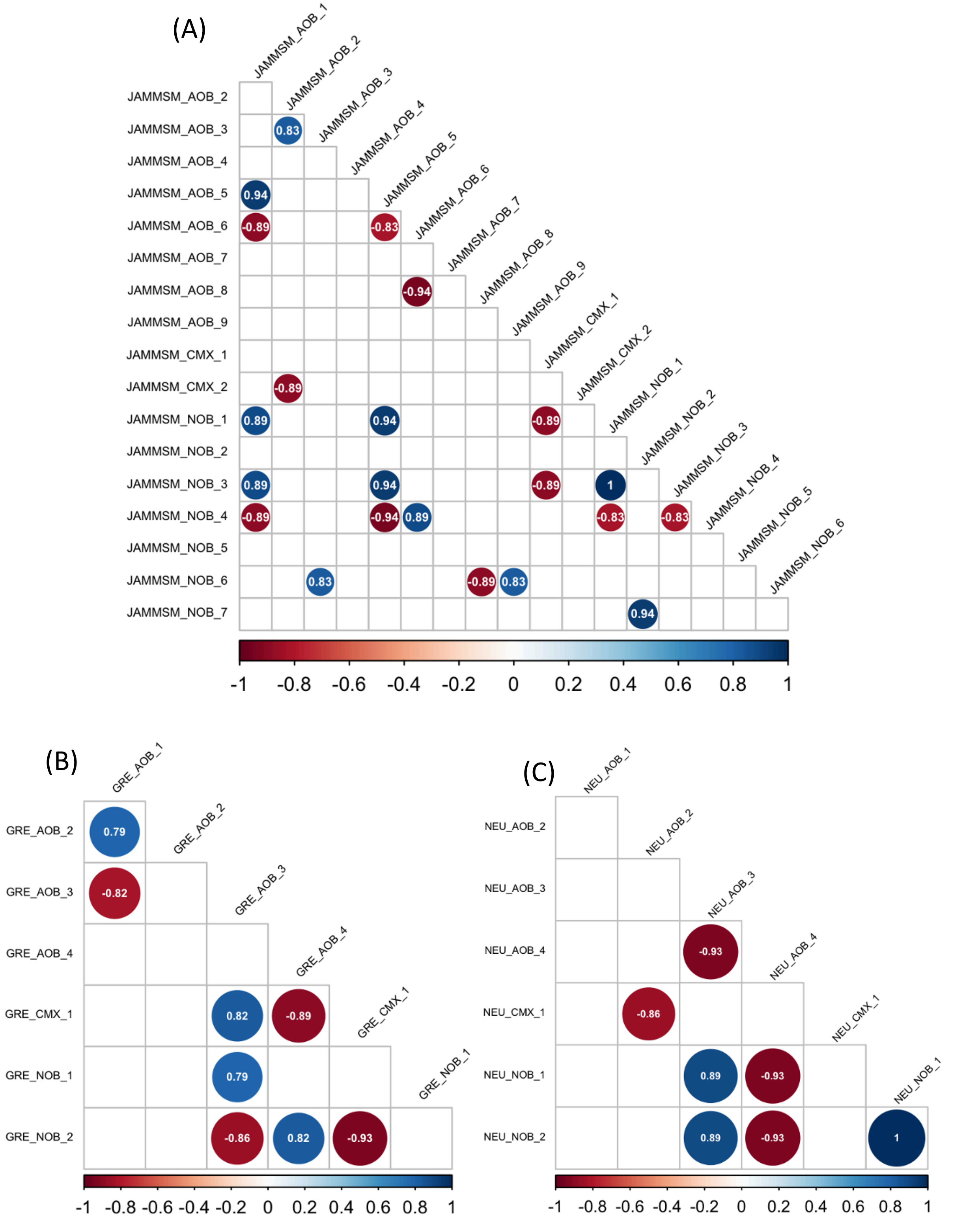


**Figure S7:** Correlation matrix between MAG abundances from (A) JAMMSM, (B) GRE and (C) NEU. Labels show the MAG names. Circles show the value of corresponding Spearman correlation coefficients. Only significant correlations are displayed. Positive correlations are showed in blue and negative correlations in red.

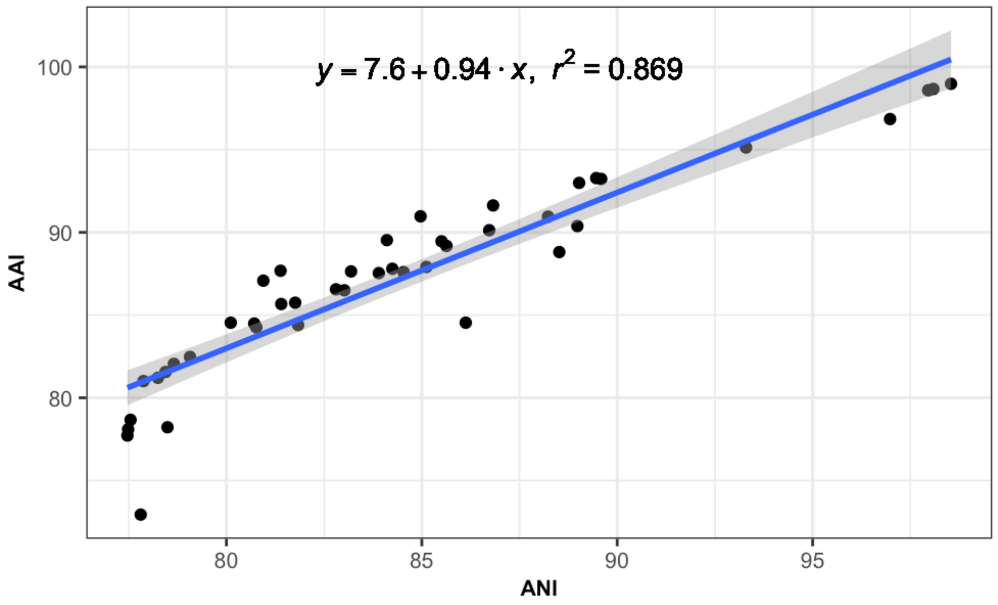


**Figure S8:** Highly positive correlation between ANI and AAI values.

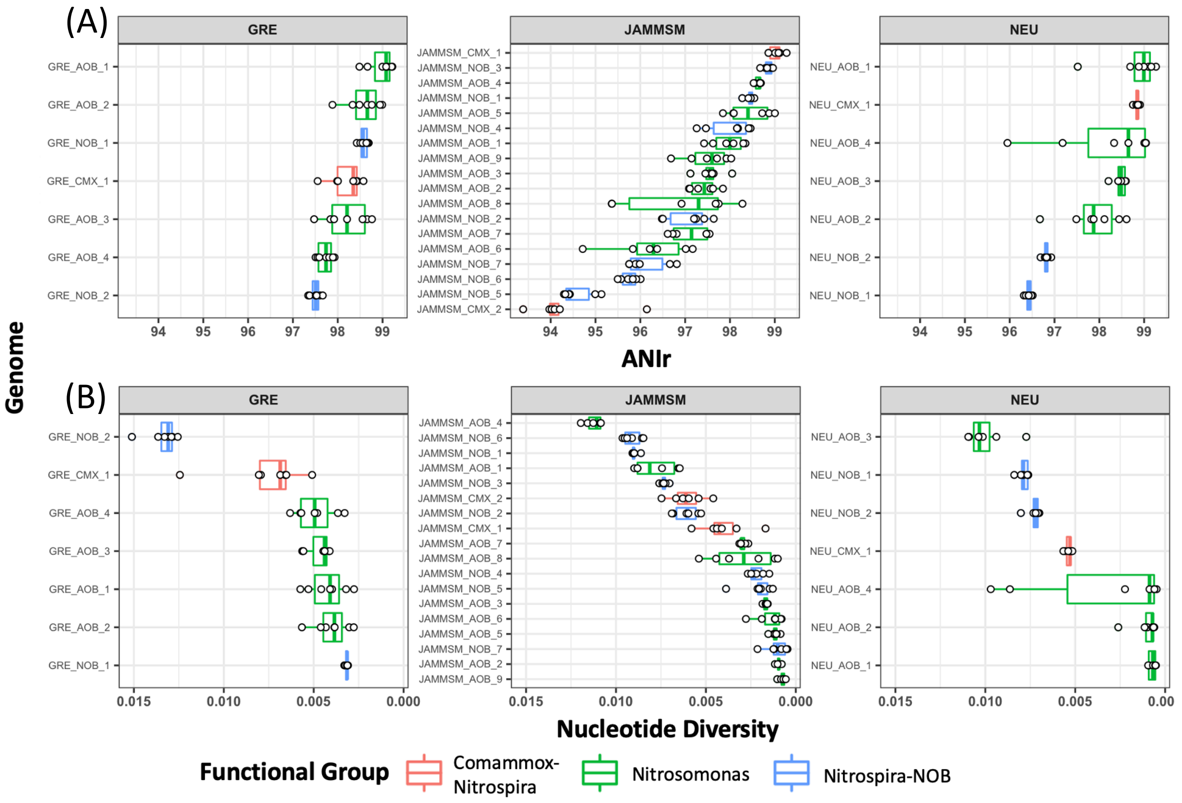


**Figure S9:** (A) ANIr and (B) nucleotide diversity values for each MAG. Points represent the average value of each MAG in one sample.

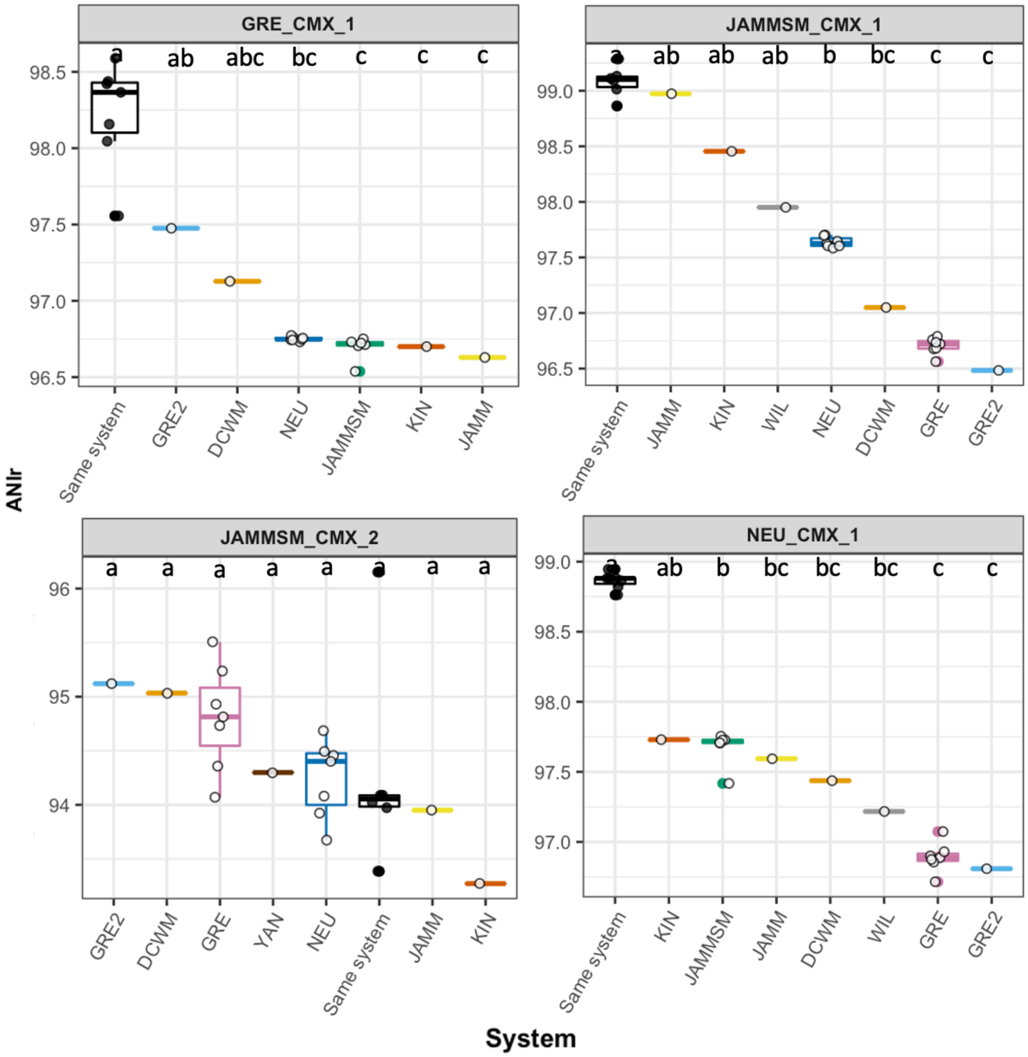


**Figure S10:** ANIr values for each comammox MAG across systems. Boxplots are colored by system and points represent the average value of each MAG in one sample of that system. Black points represent ANIr values based on mapping of metagenomes from the system where the MAG was obtained, while open circles represent ANIr values using metagenomes from the other systems.
