## Supplemental Methods for "Low diversity and microdiversity of comammox bacteria in wastewater systems suggests wastewater-specific adaptation within the *Ca*. Nitrospira nitrosa cluster"

Supplementary Methods

**Samples selection and processing**

Samples were collected monthly from the nitrifying bioreactors at GRE, NEU, and JAMMSM by operational personnel at each wastewater treatment utility and shipped overnight to Northeastern University (NU), where they were processed and stored in Lysing Matrix E tubes at -80$^{o}$C until DNA extraction. Additionally, the utilities provided environmental data and process parameters from the nitrification reactors using methods as reported previously. samples from GRE and NEU were collected from the suspended solids in the aeration zones, while samples from JAMMSM were extracted from the biofilms from integrated fixed film activated sludge (IFAS) carriers. DNA extraction was performed using the DNAeasy PowerSoil kit (Qiagen, Inc., Cat. No.12888) following manufacturer instructions with some modifications (1). Extracted DNA was quantified using Qubit 3.0 dsDNA Broad Range Assay (ThermoFisher Scientific, Cat. No. Q32850), while quality was assessed by agarose gel imaging and stored at -80$^{o}$C until further processing.

**PacBio full-length 16S rRNA gene sequencing and analysis**

The 16S rRNA gene amplicons were generated with the universal barcoded primers that amplify the full-length 16S rRNA gene (27F: AGRGTTYGATYMTGGCTCAG and 1492R: RGYTACCTTGTTACGACTT) with PCR reaction and thermocycling conditions outlined in supplementary text. in 2x Roche KAPA HiFi Hot Start Ready Mix (KAPA Biosystems) with the following thermocycling conditions: initial denaturing at 95$^{o}$C for 3 minutes, followed by 21 cycles of denaturing at 95$^{o}$C for 30 second, annealing at 57$^{o}$C for 30 seconds, and extension at 72$^{o}$C for 60 seconds. PCR products were subject to library preparation using the SMRTBell Express Template Prep kit 2.0. The resultant library was sequenced on 1 SMRTcell 8M on a PacBio Sequel IIe using the circular consensus sequencing (CCS) mode and a 10 hours movie time. Raw reads were demultiplexed using the lima application (lima --ccs --different --split-bam-named). CCS analysis was done using SMRTLink v10.0 with the following parameters: ccs --min-passes 3 --min-rq 0.999.

Downstream data processing was performed on a system-by-system basis using the DADA2 v1.19.2 (2) in R v4.0.4. Specifically, we used the DADA2 sample inference method for full-length 16S rRNA gene with single-nucleotide resolution. The DADA2 workflow included primer removal, quality filtering, dereplication, learning the dataset-specific error model, amplicon sequence variants (ASVs) inference, chimera removal, and taxonomic assignment with the SILVA SSU 138 Ref NR database using default settings (Table S1a). The three ASVs tables were rarefied with the 'rarefy_even_depth' function from the R package phyloseq v1.32.0 (3) to sample with the smallest library size per system. To compare the population composition of the systems, the three tables were combined using the 'mergeSequenceTables' function from DADA2.

ASVs were aligned with 'AlignSeqs' function of the R package DECIPHER v2.16.1. (4), the distance between sequences was calculated using the 'DistanceMatrix' function, and ASVs were clustered into operational taxonomy unit (OTUs) at 98.7% sequence identity, (5,6) with 'IdClusters'. The R package dplyr v1.0.7 (7) was used to annotate ASVs with OTU labels. Finally, 'merge_taxa_vec' function in package speedyseq v0.5.3.9018 (8) was used merge the taxonomy table with the new OTU table using the taxonomy of the most abundant ASV within each OTU as the consensus OTU taxonomy. The most abundant ASV sequence from all OTUs were aligned with MUSCLE v3.8.1551 (9), and the phylum/class phylogenetic tree was constructed and visualized using IQ-TREE v2.0.3 (10) and iTOL v2.1.7 (11), respectively. The relative abundance of each phylum/class group was calculated as the cumulative reads counts of all OTUs assigned to that group divided by total reads per sample. Principal Coordinates Analysis (PCoA) was performed with the weighted and unweighted UniFrac distance metric (12) using the 'ordinate' function of phyloseq v1.32.0 and the 'plot_ordination' function of ggplo2 v3.3.5 (13).

**Short and long read metagenomic sequencing**

The shotgun genomic libraries were prepared with the Hyper Library construction kit from Kapa Biosystems without PCR amplification and sequenced on one NovaSeq SP lane for 300 cycles (2x150nt reads) from each end of the fragments on a NovaSeq 6000. Fastq files were generated and demultiplexed with the bcl2fastq v2.20 Conversion Software (Illumina). The run resulted in a total of 1.78 billion paired-end reads (Table S1b). Six samples (two from each system), with the higher comammox bacterial abundance, were sent for long reads Oxford Nanopore sequencing (Table 1) to the Roy J. Carver Biotechnology Center at the University of Illinois Urbana-Champaign Sequencing Core. Genomic DNA from each sample were subject to Nanopore library using the SQK-LSK109 library kit. Each library was individually sequenced on a SpotON R10.3 FLO-MIN106 flow-cell for 48 hours, using a GridIONx5 sequencer, generating up to 10 Gbp of reads with an average of ~7.5 Gbp. Base-calling and adaptor trimming were performed with Guppy v3.6.1 (Table S1c).

**Phylogenetic placement of Nitrifier MAGs**

*Nitrospira* MAGs (n=15) recovered from this study were phylogenetically placed in context of 90 previously publicly available *Nitrospira* genomes downloaded from NCBI (Table S5). Prior to phylogenetic analysis, the *Nitrospira* reference genomes were dereplicated using drep v2.5.4 with default parameters resulting in 79 genomes used as references for phylogenetic tree reconstruction in Anvi'o v6.1. ORFs were predicted using Prodigal v2.6.3 and then searched against a collection of HMM models (Bacteria_71) including 38 ribosomal proteins, summarized by Lee (2019) (14) using hmmscan v3.2. (15). Multiple pairwise alignments for each gene were performed using MUSCLE v3.8.1551. The phylogenomic tree was constructed using ITOL v2.1.7. A total 64 publicly available *Nitrosomonas* genomes were downloaded from NCBI (Table S6), and after dereplication, 48 were used as reference genomes for phylogenetic tree reconstruction along with 17 *Nitrosomonas* MAGs obtained in this study. The tree reconstruction followed the same process mentioned above. Maximum likelihood phylogenetic tree of *Nitrospira*-comammox and *Nitrosomonas* based on the *haoA* and *amoA* genes, respectively, were performed after extracting each gene from MAGs and reference genomes and aligning them with MUSCLE v3.8.1551 followed by construction of the tree using IQ-TREE v2.0.3.

**Microdiversity analyses**

Average nucleotide identity from reads (ANIr) were estimated as follows. First, short reads from each sample were mapped to MAGs using BLASTn v2.9.0 to generate mappings in Tabular BLAST format (-outfmt 6 option). The BLASTn outputs were used to generate a list of hits per subject sequences using 'BlastTab.catsbj.pl'. Read recruitment plots were constructed for each MAG and sample using the 'BlastTab.recplot2.R' script from the R package enveomics.R v1.8.0 (16). Finally, ANIr values were calculated with 90% read identity threshold (the recommended threshold for intra-population) using the 'enve.recplot2.ANIr' R script from the enveomics.R package. For inStrain based analysis, MAGs from each system were concatenated into a single fasta file (i.e., one fasta file per system). Pair-end reads of each system were mapped to their respective MAGs' fasta files using Bowtie2 v2.3.5.1 (17), while gene calling was performed using Prodigal v2.6.3. Then, 'inStrain profile' was run on all resulting mapping files with default settings. The inStrain output 'genome_info.tsv' and 'scaffolds_info.tsv' were used to assess and compare genetic variation between genomes and within species. Specifically, inStrain calculates the nucleotide diversity (a measurement of genetic diversity or microdiversity) at every position along the genome using mapped reads. The nucleotide diversity was estimated in MAGs with more than 25% of bases with 5x minimum coverage (18). Finally, we ran 'inStrain compare' to calculate the population average nucleotide diversity (popANI) of the nitrifier MAGs between the samples where they were detected in a pairwise manner. Population ANI is a unique ANI calculation performed by inStrain that considers both major and minor alleles. If two populations share any alleles at a loci, including the major and minor alleles, it does not count as a difference when calculating popANI. This is different from the traditional ANI (called consensus ANI or conANI in inStrain), which only considers major alleles to call (or not) a substitution. This consideration of shared minor alleles greatly increases the accuracy of strain-level comparisons, if sufficient sequencing depth is available to confidently detect minor alleles. Based on inStrain recommendations, we used 5x as the default coverage because it is the lowest number at which minor alleles under 50% frequency can be reliably detected (18).
